# Multiplexed single-cell transcriptomic analysis of normal and impaired lung development in the mouse

**DOI:** 10.1101/868802

**Authors:** K. M. Hurskainen, I. Mižíková, D. P. Cook, C. Cyr-Depauw, F. Lesage, N. Andersson, E. Helle, L. Renesme, R.P. Jankov, M. Heikinheimo, B. C. Vanderhyden, B Thébaud

## Abstract

During late lung development alveolar and microvascular development is finalized to enable sufficient gas exchange. Impaired late lung development manifests as bronchopulmonary dysplasia (BPD) in preterm infants. Single-cell RNA sequencing (scRNA-seq) allows for assessment of complex cellular dynamics during biological processes, such as development. Here, we use MULTI-seq to generate scRNA-seq profiles of over 66,000 cells from 36 mice during normal or impaired lung development secondary to hyperoxia. We observed dynamic populations of cells, including several rare cell types and putative progenitors. Hyperoxia exposure, which mimics the BPD phenotype, alters the composition of all cellular compartments, particularly alveolar epithelium, capillary endothelium and macrophage populations. We identified several BPD-associated signatures, including Pdgfra in fibroblasts, Activin A in capillary endothelial cells, and Csf1-Csf1r and Ccl2-Ccr2 signaling in macrophages and neutrophils. Our data provides a novel single-cell view of cellular changes associated with late lung development in health and in disease.

## INTRODUCTION

Late lung development is responsible for the formation of intricate structures enabling the exchange of inspired oxygen from the atmosphere and carbon dioxide from the blood, which is the primary function of the mammalian lung. This complex task is achieved in the smallest, most distal respiratory units of the lung (the alveoli) and occurs across the alveolo-capillary barrier. The process of gas exchange takes place through an extremely thin structure of the barrier (0.2 - 2μm) and vast alveolar surface area of the lung (∼ 75m^2^). The formation of this complex structure is achieved via interconnected events of secondary septa formation and microvascular maturation during period of late lung development. These processes are facilitated by temporarily and spatially coordinated crosstalk between multiple cell types in the lung microenvironment. In addition to gas exchange, the lung acts as an important immune barrier, requiring resident alveolar macrophages to transition towards a mature anti-inflammatory phenotype. However, the signals driving these processes and the landscape of resident cells during late lung development remain largely uncharacterized^1–3^.

In humans, impaired late lung development presents as bronchopulmonary dysplasia (BPD), the most common chronic lung disease in children. BPD occurs as a consequence of premature birth, which is associated with respiratory distress and subsequent treatments in the neonatal intensive care unit^4^. In addition to impaired alveolar and microvascular formation, immune development of the lung is interrupted, leading to recurrent bacterial and viral respiratory infections. To mimic these injuries, rodent models of BPD utilize various levels of hyperoxia and/or other pro-inflammatory stimuli. Sustained exposure of neonatal mice to hyperoxia leads to a BPD-like lung phenotype, making it an ideal model to identify and study pivotal developmental steps during late lung development^5^.

Various cellular roles of late lung development have been extensively studied, establishing important functions for myofibroblasts in secondary septation, endothelial cell (EC) signaling in the processes of microvascular maturation and coordination of inflammatory cell signaling^5^. The abundance and identity of individual cell types are dynamic throughout lung development. Identification and classification of lung cells becomes even more complex under pathological conditions, in particular when the nature of the disease is heterogenous. Traditional methods to assess molecular characteristics of pathologies have depended on bulk measurements of protein or RNA, but given the heterogeneity of lung tissue and its dynamics during late development, these measurements are confounded by changes in cellular composition. As a result, changes in individual cell types cannot be identified. This is more problematic when responses are limited to rare populations, as these changes will be masked by the signal from more-abundant cell types. To circumvent these obstacles, we herein employed multiplexed single-cell RNA sequencing (scRNA-seq) to resolve changes in cellular composition and state during both normal and impaired late lung development.

Here, we report an extensive profiling of the cellular composition in the developing mouse lung by generating scRNA-seq profiles of 66,200 cells from 36 normally and aberrantly (O_2_-exposed) developing mouse lungs at three time points (P3, P7, and P14). We observed greatly diverse and dynamic populations of cells, including several rare cell types, such as distal alveolar stem cells (DASCs) or Schwann cells. Hyperoxia exposure altered the phenotype of all major cell types, particularly alveolar epithelium, capillary endothelium and macrophage populations. Furthermore, we identified multiple cell-specific gene signatures, providing a detailed cell and molecular atlas of normal and impaired post-natal lung development.

## RESULTS

### Detailed map of cellular composition during normal and impaired late lung development

In order to create a comprehensive cellular map of the normal and impaired developing lung, we generated scRNA-seq profiles of 36 mice on postnatal days (P)3, 7 and 14 (Fig. 1a-b). Arrest in lung development was induced by normobaric hyperoxia (85% O_2_) from day of birth (P0) to P14 (Fig. 1a, c). We optimized the single cell preparation protocol by testing several digestion conditions as assessed by FACS and scRNA-seq analysis (Supplementary Figure 1a-b). To evaluate the actual cell contribution *in vivo* prior to tissue digestion, we performed a stereological assessment of alveolar epithelial type 2 (AT2) cells. The number of AT2 cells *in vivo* was not impacted by hyperoxia as assessed by stereology, supporting the validity of our observations (Supplementary Figure 1c; Supplementary table 1). In total, we generated scRNA-seq expression profiles for 66,200 cells (∼11,033 cells/group) (Supplementary Figure 1d). Single-cell suspensions from individual mice were multiplexed using MULTI-seq^6^ (Supplementary fig. 1e-f). No major sex-dependent bias in cell distribution could be observed, with a single exception of a subpopulation of stromal cells (Supplementary Figure 1g-h). Cells were clustered based on their expression profile, and cell types annotated based on established cell markers available on LungMap, CellMarker, The Human Protein Atlas, and in the published literature (Fig. 2a-b, Supplementary table 2). A total of 22 clusters were identified and divided into six major cell groups: epithelial, stromal, endothelial, myeloid, lymphoid and mesothelial cells, (Fig. 2c-d, Supplementary figure 2a) which were further re-clustered (Fig. 2a, Supplementary Figure 2b). We observed dynamic changes in cellular composition of normally developing lungs, with most changes occurring between P7 and P14. Impaired alveolar development induced by hyperoxia changed the cellular distribution of the lung at all time points (Fig. 2b, Fig. 2e).

**Figure 1.**
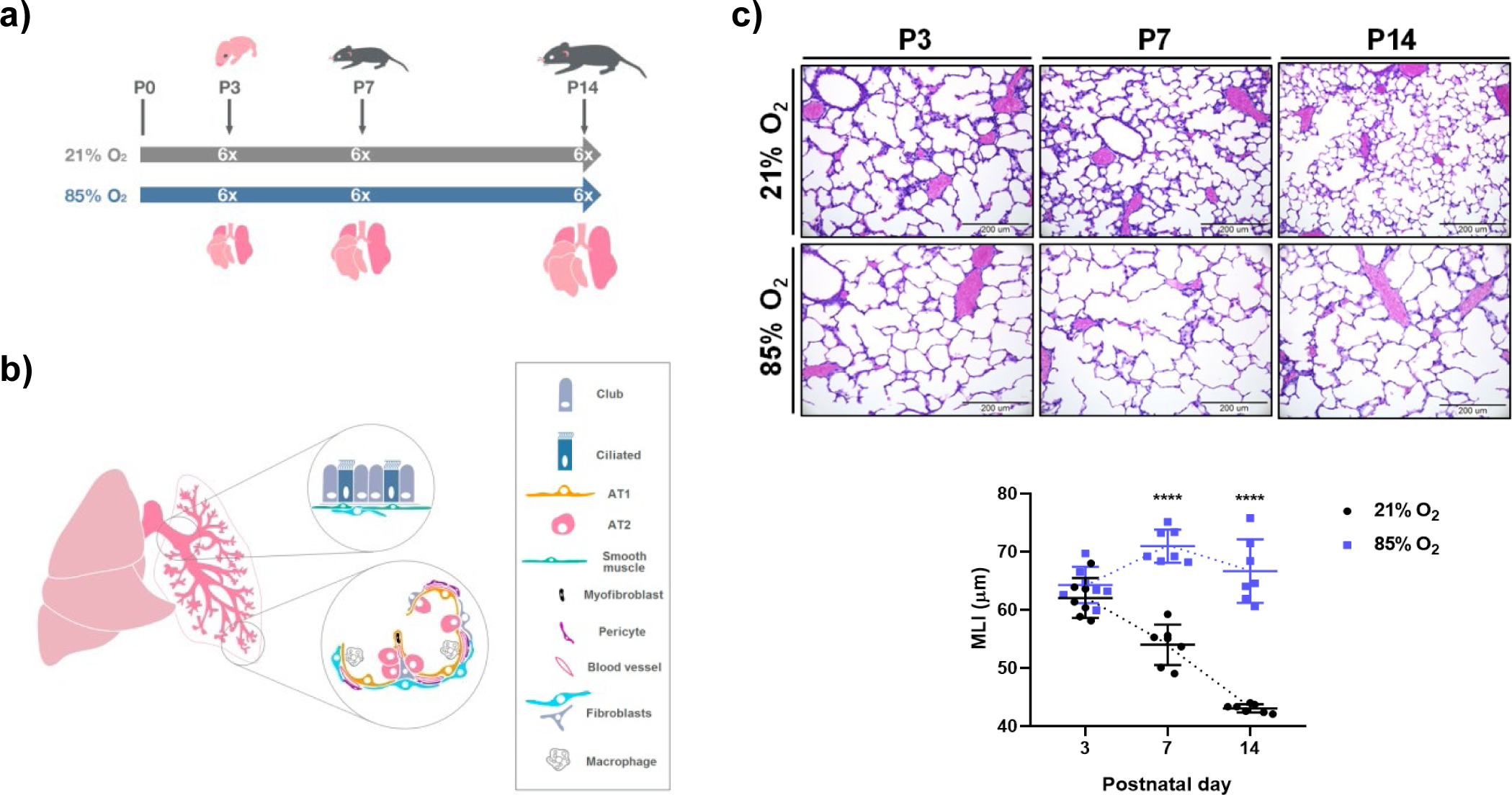
Exposure to hyperoxia induced an arrest in alveolarization in the developing mice lungs. a) Newborn mouse pups were exposed from day of birth to normal (21% O_2_) or hyperoxic (85%O_2_) environment. A total of 36 lungs were harvested on postnatal days (P)3, 7 and 14. n=6/group. b) The isolation process was optimized in order to secure an equal representation of cell types from multiple regions of the lung. c) Representative histological sections from lungs developing at normal (21% O_2_) or hyperoxic (85% O_2_) conditions at P3, P7 and P14. Average alveolar size was assessed by the mean linear intercept (MLI) measurement. Scale bar = 200µm. Data are presented as means ± SD. Statistical analyses were performed with GraphPad Prism 8.0. The presence of potential statistical outliers was determined by Grubbs’ test. Significance was evaluated by unpaired Student’s *t-*test. P values < 0.0001: ****. n = 7/group.

**Figure 2.**
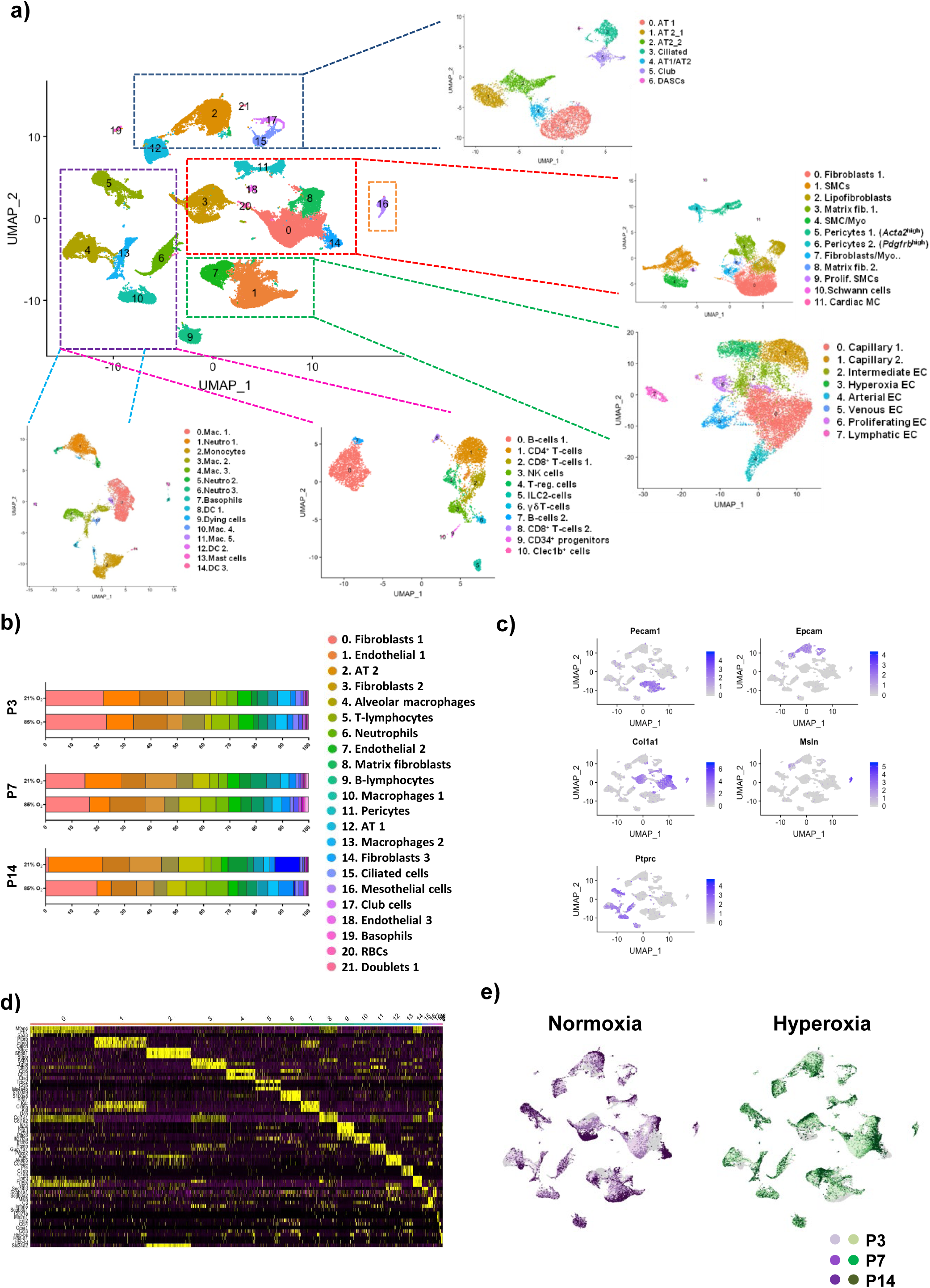
Map of cellular composition in normal and impaired late lung development. a) A total of 22 distinct cell clusters were identified and divided into 6 subsets. Epithelial, mesenchymal, endothelial, myeloid, lymphoid and mesothelial subsets were further re-clustered and analyzed separately. b) Cluster distribution in lungs of normally and aberrantly developing mice at P3, P7 and P14. n = 6/group. c) uMaps of principal identifiers of epithelial, mesenchymal, endothelial, immune and mesothelial populations. d) Heatmap of top 5 most differentially expressed genes across the 22 main clusters. e) uMaps depicting developmental trajectories in normally and aberrantly (O_2_-exposed) developing lungs as observed in real time.

### Hyperoxia alters AT2 cells population in developing lungs

We identified seven clusters of epithelial cells with distinct expression profiles (Fig. 3a-c, Supplementary Figure 3a; Supplementary table 3). Two bronchial epithelial clusters, club cells and ciliated cells, were identified. The frequency of ciliated cells in normally developing lungs remained constant between P3 and P7, but decreased significantly by P14, which was even more pronounced in hyperoxic lungs (20% *vs.* 8.8% at P7, respectively) (Fig. 3d; Supplementary Figure 3b-c). Within the alveolar epithelium, we identified one alveolar epithelial type 1 cell (AT1) cluster, two distinct AT2 clusters, and one AT1/AT2 cluster (Fig. 3a). The AT1 cluster was associated with expression of *Ager, Hopx, Akap5, and Vegfa* (Fig. 3b-c; Supplementary Figure 3a). The number of AT1 cells gradually decreased during development. This decline was more pronounced in lungs where O_2_-induced an arrest in alveolarization (Fig. 3d; Supplementary Figure 3b).

**Figure 3.**
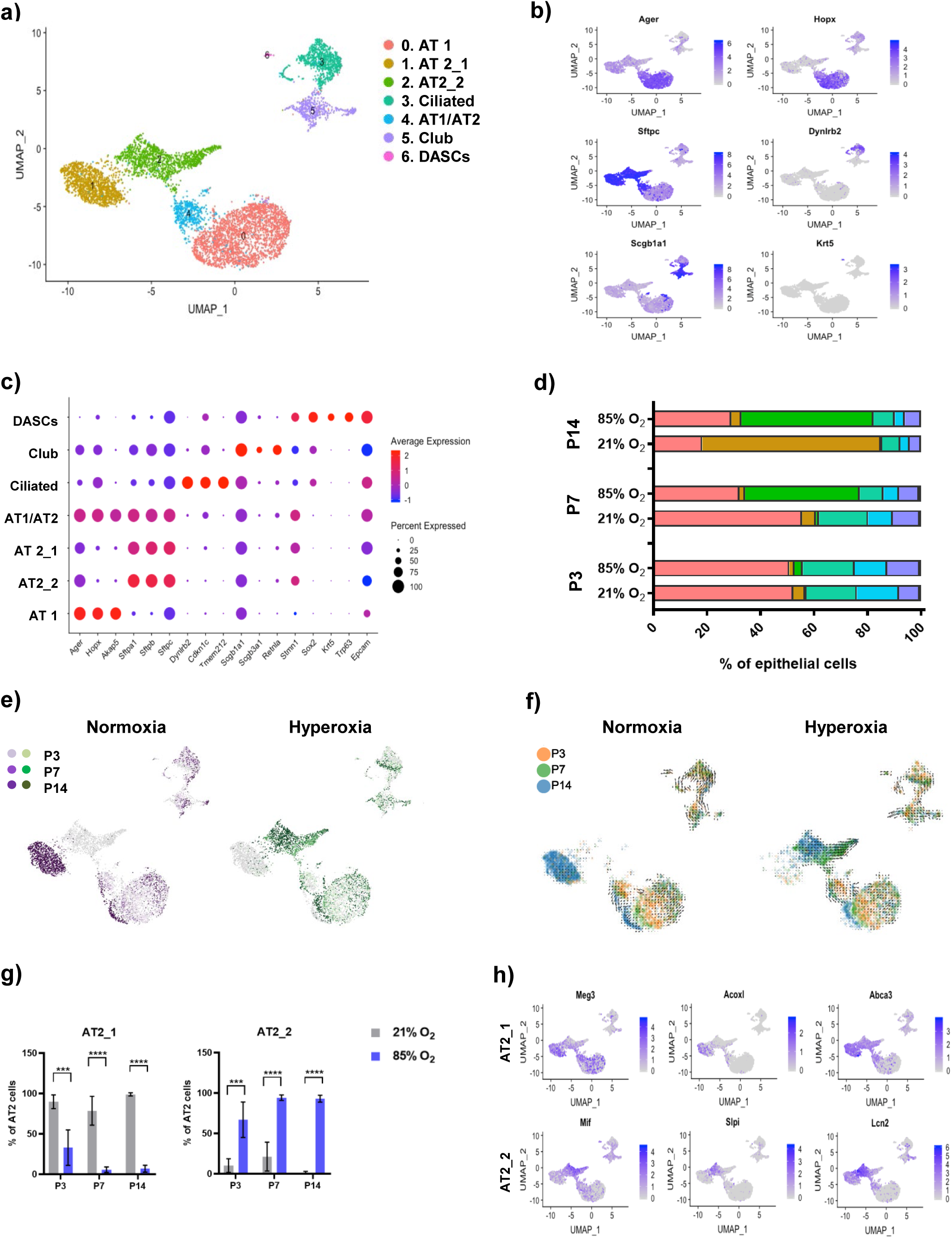
Cellular composition of epithelial cells during normal and aberrant late lung development. a) A total of 7 clusters of epithelial cells were identified in developing lungs. b) uMaps of principal identifiers of different epithelial cell types. c) Clusters were identified based on the expression of known cell type-specific markers. d) Relative contribution of individual clusters changed significantly during the development and after exposure to hyperoxia. n = 6/group. e) uMaps depicting developmental trajectories in normally and aberrantly developing lung epithelium. f) Predicted RNA velocity in normally and aberrantly developing lung epithelium. g) Hyperoxia exposure drastically altered the identity of lung AT2 cells. Data are presented as means ± SD. Statistical analyses were performed with GraphPad Prism 8.0. The presence of potential statistical outliers was determined by Grubbs’ test. Significance was evaluated by unpaired Student’s *t-*test. P values < 0.001: ***; P values < 0.0001: ****. n =6/group. h) Hyperoxia exposure altered the expression signature in developing AT2 cells. uMaps depicting expression patterns of top differentially expressed genes in clusters AT2_1 (upper panel) and AT2_2 (bottom panel).

AT2 cells had distinct transcriptional profiles in healthy and aberrant lungs (Fig. 3a-c; Supplementary Figure 3a; Supplementary tables 3-5). The AT2_1 cluster consisted almost exclusively of cells from healthy lungs at P14, while the AT2_2 cluster comprised cells from hyperoxic lungs at P7 and P14. This indicates that the AT2 population, which normally expands following the peak of secondary septation (P7), diverges in diseased lungs into a novel, undefined population (Fig. 3d-g; Supplementary Figure 3a-c). While expression levels of surfactant proteins were similar, numerous other genes were differentially expressed between the two AT2 clusters (Supplementary table 4). Among the hyperoxia-specific genes were genes coding for multiple BPD and epithelial damage-related factors, including *Slpi* - a protease inhibitor involved in defence against epithelial damage-and the innate immune response regulator *Mif* ^7^ (Fig. 3h; Supplementary Figure 3d). MIF was previously shown to promote production of IL6 and IL1β, and has been implicated in the arrest of lung development and angiogenesis^8, 9^. *Lcn-2* expression was also higher in the aberrant AT2 cells, which was shown to be associated with development of BPD^10^. Among the top normoxia-specific genes were *Meg3* and *Abca3* (Fig. 3h, Supplementary Figure 3e). *Meg3* is a known regulator of epithelial cell differentiation, while *Abca3* is a crucial factor in surfactant and lamellar bodies’ metabolism^11, 12^. Mutations of *Abca3* have been associated with interstitial lung disease and progressive respiratory distress syndrome and further, knock-down of *Abca3* resulted in abnormal lamellar bodies^13^. Supporting this notion, Gene Ontology (GO) Term analysis revealed that the surfactant homeostasis pathway is diminished in diseased lungs (Supplementary table 5). An intermediate AT1/AT2 cell cluster was apparent at all timepoints (Fig. 3a-c). Developmental dynamics and RNA velocity suggested that the AT1/AT2 cluster may represent an early form of AT2 cells and a potential source of AT1 cells (Fig. 2e and f).

In addition to major epithelial cell types, we identified a rare population of DASCs, which was positive for the DASCs-specific markers *Krt5* and *Trp63,* and negative for bronchial alveolar stem cell marker *Scgb1a1* (Fig. 3a-c). In addition, DASCs expressed *Itga6*, *Sox2*, and *Myc*, previously associated with basal epithelial cells (Supplementary Figure 3f).

We further sought to identify a subpopulation of *Axin2^+^* AT2 progenitors, postulated to play a role in alveolar regeneration^14^. We identified a small number of epithelial cells expressing *Axin2*, mostly located to AT2 clusters. Although *Axin2^+^* cells were not confined within a separate cluster, they displayed an interesting dynamic during the development as they increased with age. Notably, the *Axin2^+^* cells were further increased in the epithelium of diseased lungs at P7 by 73% compared to healthy lungs (Supplementary Figure 3g).

### Hyperoxia exposure transforms lung stromal populations

Within the stroma, we identified twelve populations, including pericytes, fibroblasts, and smooth muscle cells (SMCs) (Fig. 2a; Fig. 4a-c; Supplementary Figure. 4a; Supplementary table 6). We identified two clusters of pericytes, distinguished by expression of *Acta2* (Pericytes 1) and *Pdgfrb* (Pericytes 2) (Fig. 4a-c). All pericytes expressed pericyte-specific markers and were negative for the fibroblast marker *Pdgfra*^1, 15, 16^ (Fig. 4b-c). Both populations slowly increased during development, together representing ∼6.2% of stromal cells by P14. While *Acta2^high^* pericytes were not impacted by hyperoxia, the population of *Pdgfrb^high^* pericytes was significantly decreased at P7 and P14 (Fig. 4d; Supplementary Figure 4b-c).

**Figure 4.**
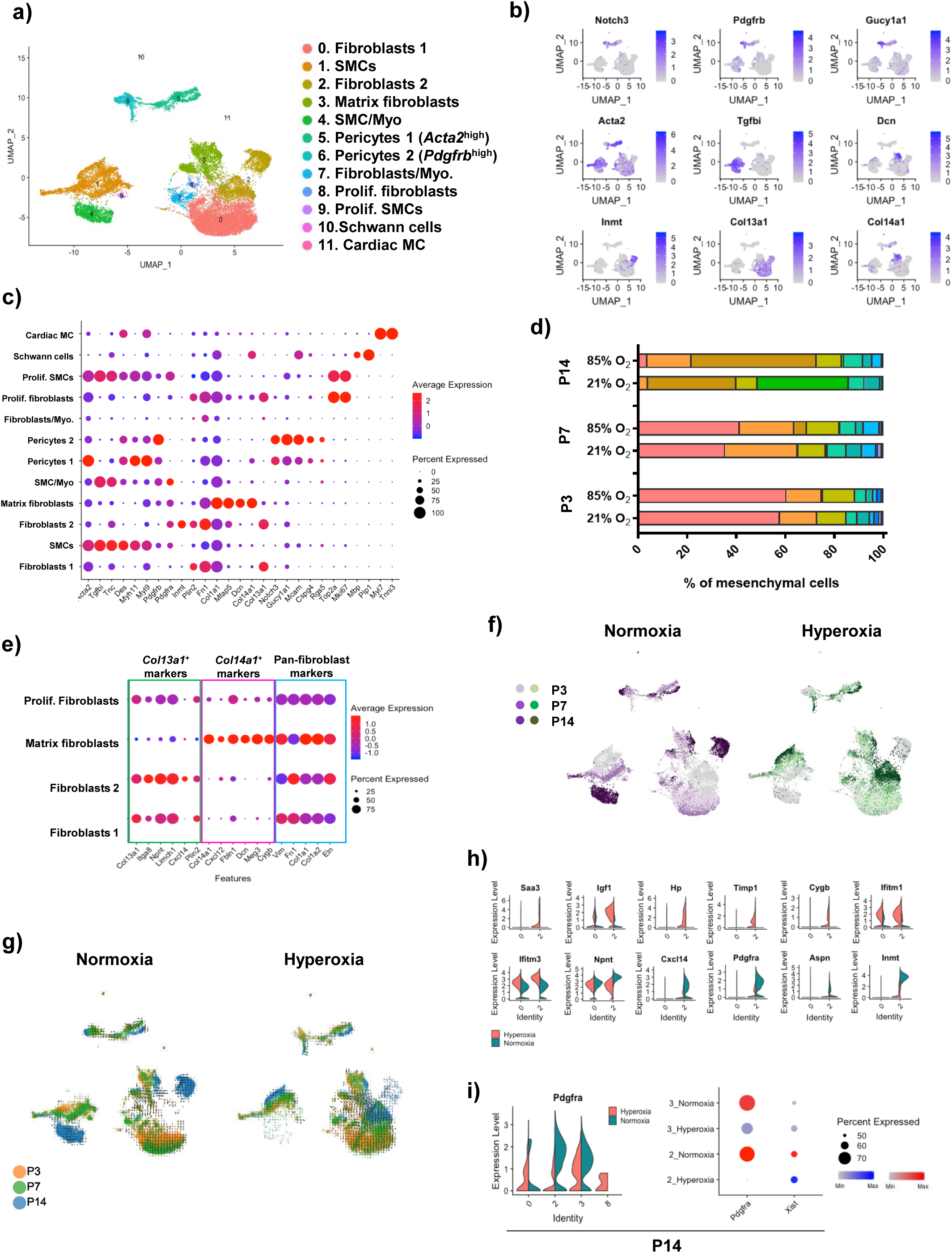
Cellular composition of mesenchymal cells during normal and aberrant late lung development. a) A total of 12 clusters of mesenchymal cells were identified in developing lungs. b) uMaps of principal identifiers of different mesenchymal cell types. c) Clusters were identified based on the expression of known cell type-specific markers. d) Relative contribution of individual clusters changed significantly during the development and after exposure to hyperoxia. n = 6/group. e) Dotplot displaying expression patterns specific to diverse fibroblasts populations. f) uMaps depicting developmental trajectories in normally and aberrantly developing lung mesenchyme. g) Predicted RNA velocity in normally and aberrantly developing lung mesenchyme h) Violin plots depicting expression of oxygen-specific markers in Fibroblast 1 and Fibroblasts 2 clusters. i) Violin plot and Dotplot depicting oxygen-specific expression of *Pdgfra* in distinct fibroblast clusters at P14.

Four distinct fibroblast clusters were identified (Fig. 4a-c; Supplementary Figure 4a; Supplementary table 6). As proposed previously^17^, we categorized fibroblasts based on the expression of *Col13a1* (Fibroblasts 1, Fibroblasts 2 and Prolif. fibroblast; clusters 0, 2 and 8, respectively) and *Col14a1* (Matrix fibroblasts, cluster 3) (Fig. 4c and e).

Among *Col13a1*^+^ fibroblasts, we identified two large (Fibroblasts 1 and Fibroblasts 2) and one smaller (Prolif. fibroblasts) population. All clusters expressed additional *Col13a1*^+^ fibroblasts markers^17^ and multiple pan-fibroblasts markers (Fig. 4e). Furthermore, all clusters expressed *Plin2*, a lipofibroblast marker. Fibroblast 1 cluster was specific to earlier timepoints in both, healthy and diseased lungs, representing ∼60% of stromal cells at P3, and almost 40% at P7, but was diminished by P14. Fibroblast 2 cells were specific to P14, representing a dominant mature fibroblast cluster (Fig. 4d). Interestingly, in hyperoxia, the Fibroblast 2 cells showed increased expression of genes found in the immature Fibroblast 1 cluster (Supplementary Figure 4d; Supplementary tables 8-9). RNA velocity and cell trajectories revealed that Fibroblasts 2 cluster likely originated from Matrix fibroblasts (cluster 3). This suggests that in hyperoxia, the mature fibroblast population was replaced by an aberrant population reminiscent of immature fibroblasts with distinct transcriptional profile (Fig. 4f-h, Supplementary table 9, 11).

The top marker for normoxic and hyperoxic parts of Fibroblast 2 cluster were *Inmt* and *Saa3*, respectively (Fig. 4h; Supplementary Figure 4e). *Saa3* has been shown to be upregulated in a lamb preterm lung injury model^18^ and to regulate Pdgfra^19^, which has a well established role in lung development in animal models^20–22^. Decreased PDGFRA expression was associated with increased risk for male patients to develop BPD^23^. The expression of *Pdgfra* was strongly decreased by hyperoxia in Fibroblasts 2 and Matrix fibroblasts at P14 (Fig. 4i; Supplementary table 9). Cells from male and female mice contributed equally to normoxic and hyperoxic portions of Fibroblasts 2 cluster (Fig. 4i). However, hyperoxic portion of Fibroblasts 2 cluster was *Pdgfra*-negative, suggesting that PDGFRA (possibly via regulation by SAA3) plays a significant role in hyperoxia-induced changes in this population.

The smallest *Col13a1*^+^ cluster (cluster 8) was characterized by expression of proliferative markers (Fig. 4a-c; Supplementary Figure 4a). Proliferative fibroblasts were specific to earlier time points in both healthy and diseased lungs, but were almost entirely absent from all lungs at P14 (Supplementary Figure 4b).

The *Col14a1^+^* cluster, Matrix fibroblasts, expressed typical markers of *Col14a1^+^* fibroblasts, including *Meg 3*, *Dcn,* and *Fbln*^17^. In addition, Matrix fibroblasts expressed *Col1a1* and *Col1a2*, consistent with the expression signature of interstitial matrix fibroblasts^1, 24^ (Fig. 4c, e; Supplementary table 6). Within the cluster, we detected a subpopulation expressing a progenitor marker *Ly6a* (*Sca1*), identifying a putative lung resident mesenchymal stromal cells (MSCs). The *Ly6a*^+^ population increased between P3 and P14 by 63% in healthy and 81% in BPD lungs. Correspondingly, gene expression levels of multiple MSCs markers (*Cd44*, *Eng* and *Thy*) were increased in cluster 3 by hyperoxia at P14 (Supplementary Figure 4f).

Among all SMCs, we identified one minor (Prolif. SMCs, cluster 9) and two major populations (SMCs and SMC/Myo, clusters 1 and 4). All clusters expressed SMC markers *Tgfbi* and *Acta2* (Fig. 4a-c; Supplementary Figure 4a). However, SMCs strongly co-expressed additional markers associated with mature muscle cells^1, 25^. Cell dynamics further revealed a developmental trajectory for SMCs cluster, gradually obtaining a more mature gene signature, characterized by expression of *Des*, *Actg2* and *Cnn1* (Figure 4f-g; Supplementary Figure 4g-h). While SMCs cluster was present at all timepoints in all lungs, SMC/Myo cluster was exclusive to healthy lungs at P14 (Fig. 4f-g; Supplementary Figure 4b-c). Cells belonging to this normoxia-specific cluster largely shared the expression signature of SMCs cluster at P14, but lacked the expression of maturity-associated genes. Instead, SMC/Myo cluster exhibited a myofibroblasts-associated expression signature^17^ (Supplementary Figure 4g). GO Term analysis confirmed the activation of important developmental pathways in SMC/Myo cluster, including fibroblast proliferation and PDGFR signaling (Supplementary table 12). The Prolif. SMCs cluster, characterized by expression of proliferation markers, was specific to earlier time points in healthy lungs, while almost entirely absent from diseased lungs at all time points (Fig. 4a-c; Supplementary Figure 4a-b).

Finally, we identified an additional population (Fibroblasts/Myo.), with an unusual gene expression pattern (Fig. 4a-c; Supplementary Figure 4a). This cluster was negative for expression of *Col13a1*, *Col14a1*, but expressed a mixture of fibroblasts, myofibroblasts and SMCs markers (Supplementary Figure 4i).

### Hyperoxia dramatically alters capillary endothelial cell development

We identified eight distinct EC clusters based on their expression profiles (Figure 5a-c, Supplementary Figure 5a, Supplementary table 13). Seven of the clusters were found in normally developing lungs, whereas one of the clusters (cluster 3) was specific to hyperoxia-exposed lungs (Figure 5a-c; Supplementary Figure 5b-c). We classified clusters 0, 1, 2 and 3 as capillary endothelial cells (Capillary 1, Capillary 2, Intermediate ECs, Hyperoxia ECs)^26, 27^ (Fig. 5a-c, Supplementary Figure 5d). Throughout normal development, the proportion of Capillary 1 cells increased significantly (Supplementary Figure 5b, e). During maturation of these cells, we observed up-regulation of several genes involved in pathways such as endothelial cell migration and angiogenesis (Supplementary Figure 5f). In contrast to the Capillary 1 cluster, the number of Capillary 2 cells decreased with time, indicating the importance of this population in early postnatal development (Supplementary Figure 5b, e). Cluster 2 was identified as Intermediate capillary ECs between clusters 0 and 1, representing less differentiated capillary ECs (Fig. 5d).

**Figure 5.**
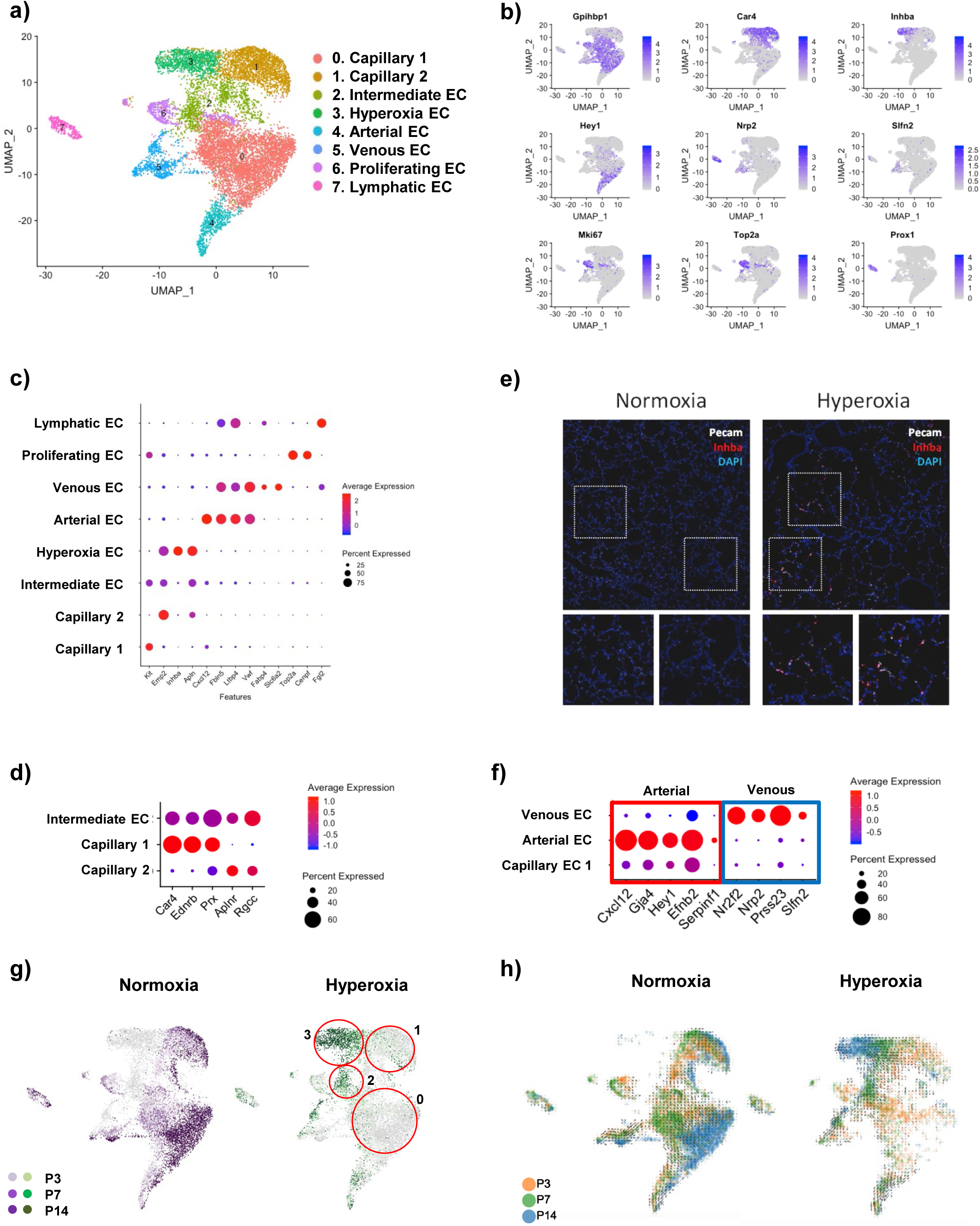
Cellular composition of lung endothelium during normal and aberrant late lung development. a) A total of 8 clusters of endothelial cells were identified in developing lungs. b) uMaps of principal identifiers of different types of endothelial cells. c) Clusters were identified based on the expression of known cell type-specific markers. d) Dotplot displaying expression patterns specific to diverse populations of capillary endothelium. e) Representative images from fluorescent RNA *in situ* hybridization for hyperoxia-specific marker *Inhba* (red) and pan-endothelial marker *Pecam* (white). Magnification: 40x. f) Dotplot displaying expression patterns specific to venous and arterial endothelial populations. g) uMaps depicting developmental trajectories in normally and aberrantly developing lung endothelium. h) Predicted RNA velocity in normally and aberrantly developing lung endothelium.

Hyperoxia significantly reduced the number of both Capillary 1 and 2 cells, and introduced a new cluster of capillary 2-like cells, Hyperoxia ECs (Cluster 3), the size of which increased with time (Fig. 5a, Supplementary Figure 5b, g). The top marker for Hyperoxia ECs was *Inhba* (Fig. 5b-c, Fig. 5e), a member of the TGFβ superfamily suggested to contribute to the pathology of BPD^28^. Several genes known to be induced by cellular stress or inflammation^29–31^ were up-regulated in the Hyperoxia EC cluster (Supplementary Figure 5h). We observed increased expression of *Ctgf* and *Fxyd5*, known to contribute to inflammatory lung injury^32, 33^ and of *Tgfb2,* shown to be associated with profibrotic responses in the lung^34^. In addition, an anti-angiogenic gene expression profile was observed, characterized by a decrease in expression of *Vegfa*, and an increase in expression of *Cdkn1a, Timp3*, *Serpine1,* and *Igfbp7* (Supplementary Figure 5h), consistent with the crucial role of angiogenic growth factors in normal lung development^35^. Further, we observed upregulation of angiogenic growth factor *Angpt2*, which mediated epithelial necrosis and pulmonary edema in hyperoxic acute lung injury^36^. Consistent with previous reports ^29^, the cell cycle inhibitor *Cdkn1a,* which may protect the lung from oxidative stress^37, 38^, was overexpressed in hyperoxia. Another potentially protective gene that was overexpressed in hyperoxia was Apelin, which was shown to reduce potential pulmonary inflammation, fibrin deposition, and partially restore alveolarization in rat pups with neonatal hyperoxic lung injury^39^.

In contrast to capillary endothelial cells, hyperoxia caused only minor changes in the arterial, venous, and lymphatic ECs (clusters 4, 5, and 7, respectively). As described previously, the expression of arterial-specific genes formed a continuum with the arterial gene expression of Capillary 1 cluster, whereas in the Venous EC cluster, the expression of arterial-specific genes was absent or considerably down-regulated (Fig. 5f).

The EC progenitors in the lung are not well understood^40^. Based on the developmental trajectory analysis we identified a putative early capillary progenitor cluster of Proliferating ECs (cluster 6) (Fig. 5c; Supplementary Figure 5a)^41^. The Proliferating ECs, together with the Capillary 1 and Intermediate capillary EC clusters, expressed *Kit*, a previously suggested endothelial progenitor marker^42^ (Supplementary Figure 5i). Hyperoxia significantly reduced the number of Proliferating ECs and expression of *Kit* at P7, a crucial stage of capillary development in the lung. In contrast to *Kit*, the expression of *Bst1*^43^ and *Procr*^44^ genes, suggested to be specifically expressed by EC progenitors, was up-regulated in the lungs by hyperoxia (Supplementary Figure 5j). However, the developmental cell trajectory analysis did not directly support the notion of progenitor nature of these cells (Fig. 5g-h).

### Hyperoxia exposure alters myeloid cell distribution, and leads to the emergence of new populations of activated macrophages and neutrophils

Immune cell clusters were identified^1, 2, 45, 46^ (Fig. 2a) and grouped as belonging to either myeloid or lymphoid lineage. During normal lung development, the relative proportion of lymphoid cells from total lung cells increased with time, representing the major immune cell type of the lung at P14 (Fig. 6a). In hyperoxia, the myeloid cells remained the most abundant immune cell type in the lung during development.

**Figure 6.**
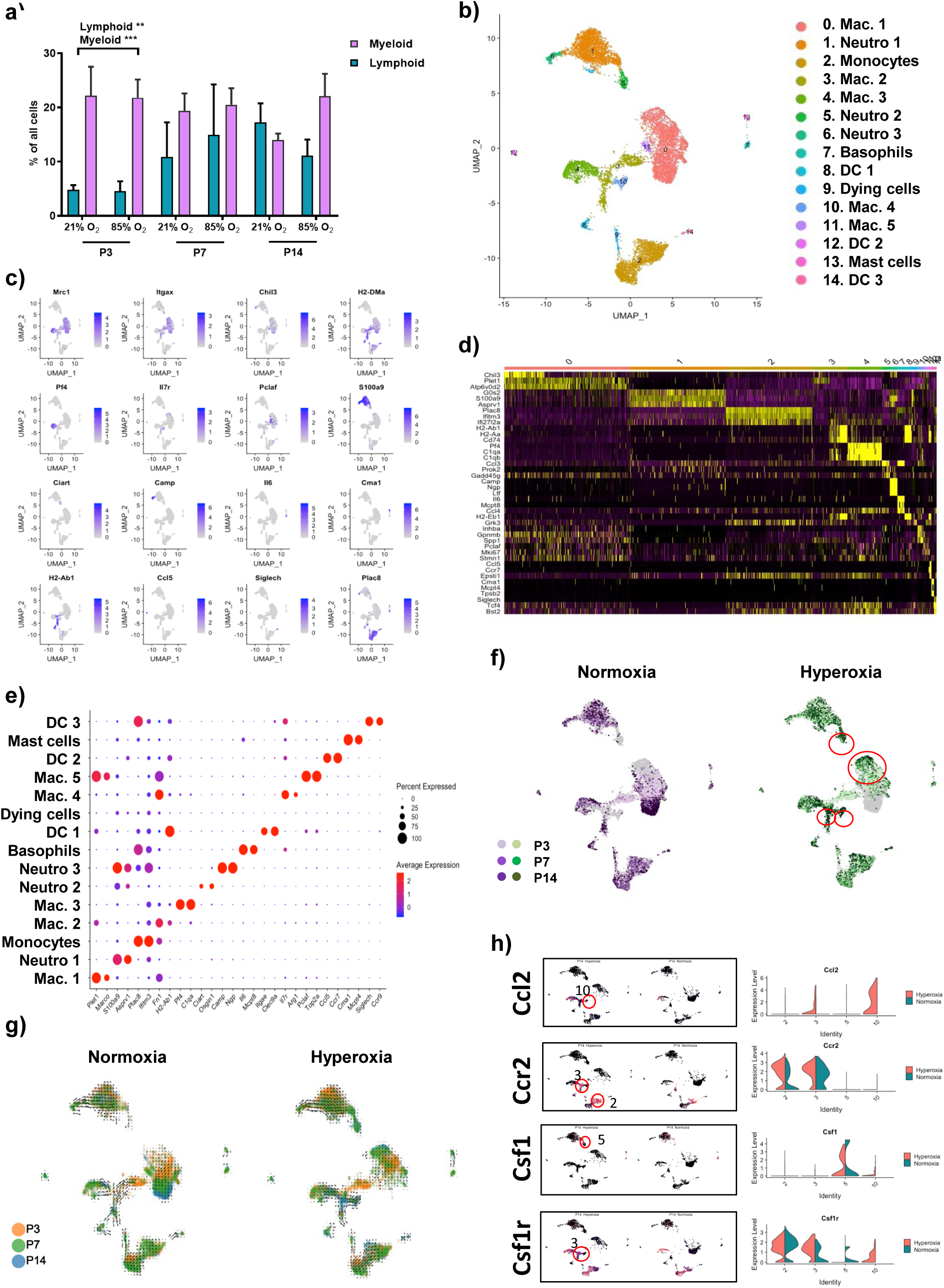
Cellular composition of lung myeloid populations during normal and aberrant late lung development. a) The relative proportion of myeloid and lymphoid cells in developing lungs was significantly impacted by hyperoxia exposure. Data are presented as means ± SD. Statistical analyses were performed with GraphPad Prism 8.0. The presence of potential statistical outliers was determined by Grubbs’ test. Significance was evaluated by unpaired Student’s *t-*test. P values < 0.01: **; P values < 0.001: ***. n =6/group. b) A total of 15 clusters of myeloid cells were identified in developing lungs. c) uMaps of principal identifiers of different types of myeloid cells. d) Heatmap of top 3 most differentially expressed genes across the 15 myeloid clusters. e) Clusters were identified based on the expression of known cell type-specific markers. f) uMaps depicting developmental trajectories of myeloid populations in normally and aberrantly developing lungs. g) Predicted RNA velocity of myeloid populations in normally and aberrantly developing lungs. h) Cell cluster and condition-specific expression patterns of *Ccr2* and *Csf1r* and their ligands *Ccl2* and *Csf1*.

We identified 15 distinct clusters of myeloid cells, including various populations of macrophages, monocytes, neutrophils, dendritic cells (DCs), mast cells and basophils (Fig. 6b-e). Hyperoxia dramatically changed the cell distribution and developmental trajectories, particularly in the macrophage populations (Fig. 6f-g, Supplementary Figure 6a-c).

Further, we identified five distinct macrophage clusters (0, 3, 4, 10 and 11) (Fig. 6b; Supplementary Figure 6d). Macrophage clusters were identified as alveolar (0, 3, 10 and 11) or interstitial (cluster 4) based on the expression of *Lgals3*, *C1qb* and *Adgre1* gene^47^ (Supplementary Figure 6e). We identified one normoxia-specific cluster, Mac.5 (Cluster 11), one partially hyperoxia-specific cluster, Mac.2 (Cluster 3), and one completely hyperoxia-specific macrophage cluster, Mac 4 (Cluster 10). This is in agreement with previously published macrophage clusters by *Mould et al*^45^.

In hyperoxia, we observed a significant decrease in the size of the major alveolar macrophage cluster, Mac.1 (Supplementary Figure 6b). Simultaneously, differential gene expression between normoxia and hyperoxia suggested a transcriptional shift characterized by down-regulation of RNA transcription and B-cell differentiation genes, and up-regulation of genes involved in chemotaxis, smooth muscle cell proliferation, inflammatory response, phagocytosis, and oxidative stress (Fig. 6f; Supplementary Figure 6f-g). Homeostatic alveolar macrophages in this cluster were characterized by the expression of *Ear1*, whereas the hyperoxic macrophages lacked the expression of *Ear1*, but showed an increase in the expression of *Marco* (Supplementary Figure 6h, i).

Alveolar macrophages were previously suggested to be self-renewing^48^. *Mould et al.* postulated that the proliferating alveolar macrophages were responsible for self-renewal of alveolar macrophages in their study. The Mac.5 cluster expressed several genes suggesting active proliferation such as *Mki67, Pclaf* and *Top2a* and was significantly diminished by hyperoxia (Supplementary Figure 6b). Our results suggest that hyperoxia exposure reduced the number of homeostatic alveolar macrophages and almost entirely diminished the proliferating, self-renewing alveolar macrophage population.

The recruitment of new monocytes/macrophages to the alveolar niche is tightly regulated^49^, allowing new macrophages to enter only if the niche is available. Consistent with this concept, we observed a reduction in the alveolar macrophage count during hyperoxia exposure, and subsequent appearance of two new hyperoxia-specific macrophage populations, clusters Mac.2 and Mac.4 (Supplementary table 14; Fig. 6b; Supplementary Figure 6d). Differential gene expression in the Mac.2 population between healthy and diseased lungs suggested that the Mac.2 cells were actively involved in innate immune response, inflammation, antigen presentation via MHC class I and organ regeneration. Concurrently, genes involved in processes such as antigen presentation via MHC class II, and regulation of T-cell differentiation were down-regulated in Mac.2 cells after hyperoxia exposure (Supplementary Figure 6j). Another macrophage cluster, Mac.4 (cluster 10), was found to be entirely hyperoxia-specific (Fig. 6f; Supplementary Figure 6b). Interestingly, we observed up-regulation of *Csf1r* and *Ccr2* in Mac.2 population after hyperoxia exposure. Furthermore, the expression of their ligands, *Csf1* and *Ccl2*, was up-regulated in the hyperoxia-specific clusters Neutro.2 and Mac.4, respectively (Fig. 6h).

These observations are consistent with the study by *Kalymbetova et al*, where *Ccr2*^-/-^ and CSF1R-depleted mice developed milder structural changes in hyperoxia-exposed lungs^50^. The cell count and transcriptional activity of Mac.3 cells (cluster 4), was stable in normal development, as well as after exposure to hyperoxia (Supplementary Figure 6a-c), and expressed high levels of inflammatory mediators, consistent with data by *Gibbings et al.*^51^ (Supplementary Figure 6k).

The concept of M1/M2 polarization of macrophages has been linked to normal development, as well as to several lung pathologies, including BPD^2, 52, 53^. Consistent with previous studies, our results suggested that the expression of M2 genes was increased in alveolar macrophages during postnatal development^2^ (Supplementary Figure 6l). Furthermore, hyperoxia enhanced the M1 signature of Mac.2 macrophages and induced a new population of macrophages - Mac.4 - characterized by the activation of both M1 and M2 genes (Supplementary Figure 6m).

Monocytes are commonly identified as functionally distinct groups of classical or non-classical monocytes based on the expression of *Ly6C*, *Ccr2,* and *Cx3cr1*. In our study, we found a single monocyte cluster, part of which was positive for classical, and part for non-classical monocyte markers. The number of classical monocytes defined by the expression of *Ly6c2* increased after hyperoxia exposure, whereas the number of non-classical monocytes defined by the expression of *Fcgrn* was reduced (Supplementary Figure 6n). These results suggest that, in addition to the hyperoxia-induced upregulation of the *Ccl2*-*Ccr2* axis in macrophages, also the number of *Ccr2* expressing classical inflammatory monocytes was increased.

We identified three neutrophil clusters: Neutro.1-3 (clusters 1, 5 and 6) (Supplementary Figure 6o). Exposure to hyperoxia caused a considerable increase in the size of the Neutro.1 cluster (Fig. 6f; Supplementary Figure 6a-b). However, hyperoxia did not induce significant changes in gene expression in these cells (Fig. 6f and g, Supplementary Figure 6c). Neutro.2 cells were hyperoxia-specific and their frequency increased with age in the lungs exposed to hyperoxia (Supplementary Figure 6a-c). Differential expression pattern in this cluster suggested decreased transcriptional activity and up-regulation of chemotactic properties (Supplementary Figure 6p). The size and transcriptional profile of the Neutro.3 cluster were unaltered during normal development and by hyperoxia (Supplementary Figure 6a-c). The developmental trajectory, inferred both by RNA velocity and time point analysis, suggested that cells in the Neutro.1 cluster originated from cells in Neutro.3 cluster, whereas Neutro.2 population seemed to consist of newly recruited neutrophils as a response to hyperoxia-induced injury (Fig.6f-g).

Several small myeloid cell clusters with a specific gene signature were also identified, including basophils, mast cells, three distinct DC populations (DC1-3)^46^ and a small cluster of dying cells (cluster 9) (Supplementary table 14; Supplementary Figure. 6q-r).

### Normal development of adaptive immunity in the lung is disturbed by hyperoxia

Based on known cell markers, we identified 11 lymphoid cells clusters (Fig. 7a-d; Supplementary table 15), including B cells, T cells, NK cells and ILC2 (Fig. 7e-f). Hyperoxia exposure considerably decreased the proportion of B-cells and CD4^+^ T-cells in the lung at P14 (Fig. 7e). Hyperoxia did not induce expression changes in lymphoid populations (Fig. 7g), whereas genes down-regulated by hyperoxia in alveolar macrophages (Mac.1) were involved in the differentiation of B-cells (Supplementary Figure 6g). Furthermore, the Mac.2 cells down-regulated pathways involved in T-cell differentiation and antigen presentation through MHCII (Supplementary Figure 6j) under hyperoxia condition. Our results suggest that hyperoxia altered the normal development of adaptive lung immunity, particularly a reduction in B-cell and CD4^+^ T-cell immunity, resulting in the myeloid cells remaining the major immune cell population of the lung at P14.

**Figure 7.**
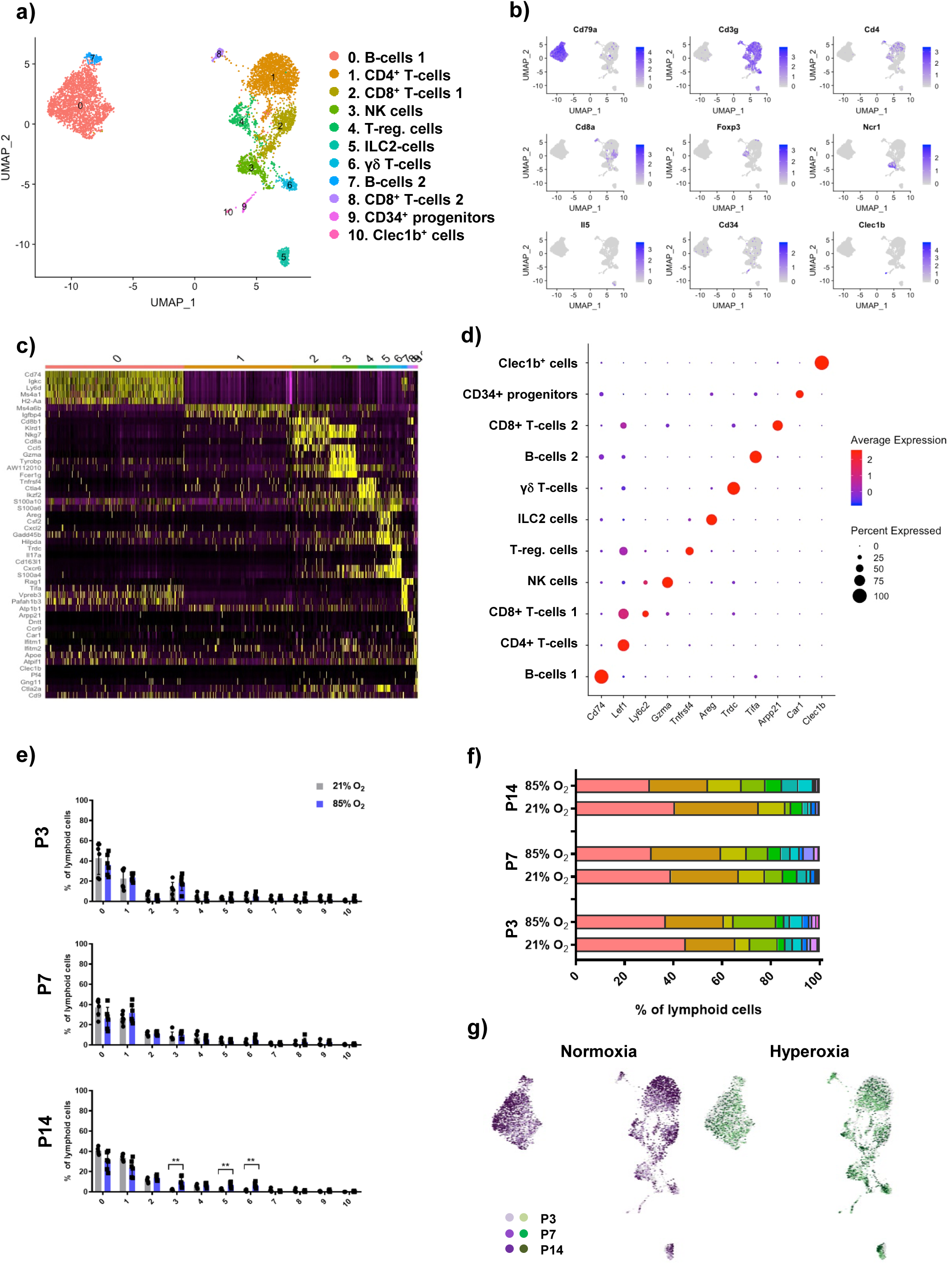
Cellular composition of lung lymphoid populations during normal and aberrant late lung development. a) A total of 11 clusters of lymphoid cells were identified in developing lungs. b) uMaps of principal identifiers of identified types of lymphoid cells. c) Heatmap of top 5 most differentially expressed genes across the 11 lymphoid clusters. d) Clusters were identified based on the expression of known cell type-specific markers. e) Hyperoxia exposure altered the relative contribution of distinct lymphoid clusters at all investigated timepoints. Data are presented as means ± SD. Statistical analyses were performed with GraphPad Prism 8.0. The presence of potential statistical outliers was determined by Grubbs’ test. Significance was evaluated by unpaired Student’s *t-*test. P values < 0.01: **. n =6/group. f) Relative contribution of individual lymphoid clusters changed significantly during the development and after exposure to hyperoxia. n = 6/group. g) uMaps depicting developmental trajectories in normally and aberrantly developing lung lymphoid populations.

### Hyperoxia alters the expression profile of mesothelial cells

A single mesothelial cluster (Fig. 2a and c) was identified based on the expression of *Msln* and other previously published markers^24^ (Fig. 8a, Supplementary table 16). Although mesothelial cells represented only 1.27% of all cells, a clear pattern of gradual decrease was observed in healthy lungs between P3 and P7, which was absent in hyperoxic lungs (Supplementary Figure 2e). The developmental arrest was associated with multiple changes in gene expression and signalling pathways (Fig. 8b-d; Supplementary tables 16-17). Among the most up-regulated were multiple anti-angiogenic factors including *Angptl7*, *Hif1a* and *Igfbp6*. Notable was the gradual increase in expression of *Timp1* and *Timp3*, as well as *Timp3* regulator *Foxf1*. FOXF1 induces fetal mesenchyme proliferation and its de-regulation was associated with lung immaturity and lung vascular defects^42, 54^. Increased *Timp1* expression was reported in ventilated preterm human lungs and murine BPD models^55, 56^, while an increase in *Timp3* expression was associated with BPD severity^57^.

**Figure 8.**
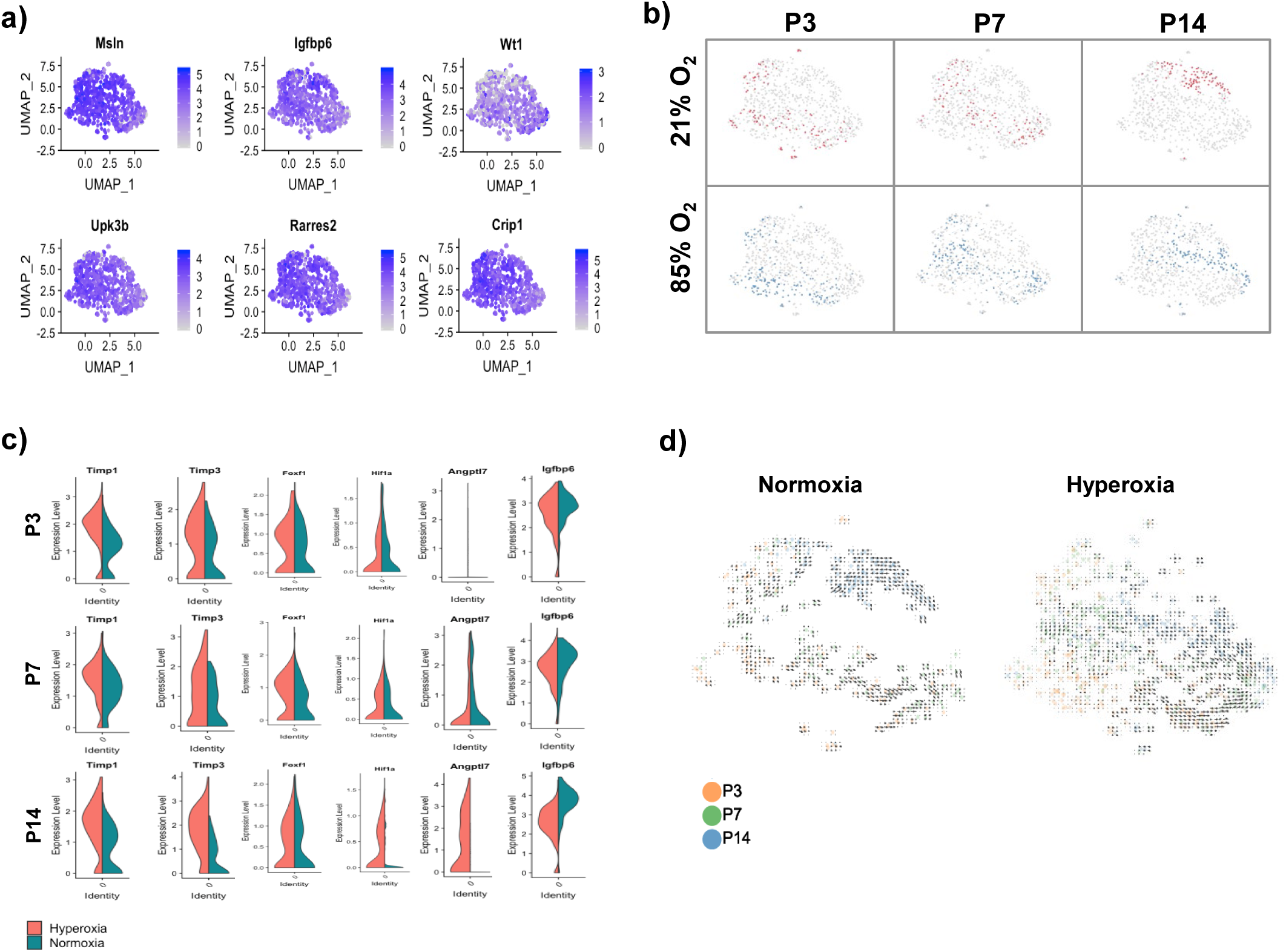
Cellular composition of lung mesothelium during normal and aberrant late lung development. a) uMaps of principal identifiers of mesothelial cells in developing lung. b) uMaps representing the temporal changes in gene expression in mesothelial population during normal and aberrant late lung development. c) Hyperoxia exposure altered gene expression in developing lung mesothelium at all investigated timepoints. d) Predicted RNA velocity in normally and aberrantly developing lung mesothelium.

## DISCUSSION

Heterogeneity and origin of lung cell lineages in the context of lung development have been the focus of significant research efforts during the last decades. Here we provide an extensive profiling of cellular composition in normal and impaired late lung development using multiplexed scRNA-seq to assess the expression profile of 66,200 cells during normal and impaired late lung development. Our study provides novel insight into the pathogenesis of impaired alveolarization by characterizing several pathological cell populations with distinct molecular expression profiles and, in so doing, highlights new putative drug targets for therapeutic interventions in BPD.

We followed the developing lung through three crucial time points of late lung development, across which we identified 7 epithelial, 12 stromal, 8 endothelial, 15 myeloid, 11 lymphoid and 1 mesothelial cell clusters. Cluster annotations were largely consistent with previously published data^1, 2, 58^. By assessing tissues at multiple time points and inferring cell dynamics using RNA velocity, we captured developmental trajectories of these populations during normal and impaired development.

In addition to major cell types, we identified several rare, putative progenitor populations. Within epithelial cells, we identified a small, but well defined population of DASCs. Within the stromal cell types, a population of lung resident MSCs was more abundant during later timepoints and was considerably impacted in developing lungs by hyperoxia. A population of endothelial cells, identified as putative capillary progenitors, was evident, particularly at early timepoints, reflecting the temporal regulation of lung development. Finally, a putative proliferating population responsible for self-renewal of alveolar macrophages was also identified. Numerous other markers that were previously proposed as progenitor-defining were expressed within particular clusters; however none defined an isolated subpopulation. Furthermore, RNA velocity and cell trajectory analyses did not provide any evidence of their progenitor properties, underscoring the need for further functional and lineage-tracing studies.

Importantly, exposure to hyperoxia impaired the composition and expression patterns of all cellular compartments necessary for normal alveolarization. Multiple affected transcriptional programs were related to cell proliferation, gene transcription, or protein translation. Within the epithelium, a population of AT2 cells, which normally expanded after the peak of secondary septation, was comprised in hyperoxic lungs by an entirely distinct aberrant population of cells, characterized by disturbed patterns of gene expression and regulatory pathways involved in surfactant homeostasis.

Within the stroma, fibroblasts found in diseased lungs failed to mature and instead, showed a less-mature expression profile normally present at earlier developmental stages. In addition, strong downregulation of *Pdgfra*, a known regulator of alveolar formation, was observed in the aberrant fibroblast population.

Particularly striking was an extensive loss of endothelial cells in diseased lungs. A putative endothelial progenitor population was almost entirely abolished by hyperoxia, and simultaneously, the number of capillary endothelial cells was significantly diminished, suggesting that hyperoxia impairs endothelial progenitors at a critical stage of capillary development. Our data supports previous studies showing that capillaries are most vulnerable to hyperoxic conditions^40, 59–61^. Furthermore, the remaining capillary endothelial cells had a pathological gene expression under hyperoxia characterized by upregulation of *Inhba* and many pro-inflammatory and anti-angiogenic markers. Specifically, our results suggest that hyperoxia induces *Inhba* expression in capillary endothelial cells indicating a significant, unexplored role for Activin A in impaired alveolar vasculogenesis.

Our study highlights the importance of the innate immune response in impaired lung development. We showed that in homeostasis, lymphoid cells expand during late lung development, becoming the prevalent immune cell type of the lung, whereas under hyperoxic conditions, myeloid cells remained the major immune cell type. The role of immune cell signaling, such as IL-1b^62^, IFNγ^63^ and TGFβ^64^, as well as the NRLP3 inflammasome^65^, has been investigated in impaired lung development by several studies. Recently, recruited *Csf1r*^+^ and *Ccr2*^+^ macrophages were shown to play pivotal roles in aberrant development, where *Ccr2*^-/-^ mice developed only moderate, and CSF1R-depleted mice developed no structural changes in lung when exposed to hyperoxia for 10 days^50^. In this study, we observed an up-regulation of *Csf1r* and *Ccr2* in hyperoxia-induced macrophages and increased *Ccr2* expression in monocytes. Moreover, the expression of *Csf1r* and *Ccr2* ligands was also increased in hyperoxia-specific neutrophil and macrophage clusters. Consistent with previous findings, our data show that hyperoxia alters lung macrophage populations through CSF1-CSF1R and CCL2-CCR2 axes.

Here we provide a detailed cell map of normal and aberrant late lung development associated with murine BPD-like lung injury. By careful optimization we limited the population distribution bias and successfully identified even some rare cell populations, including DASCs and Schwann cells. We have demonstrated extensive changes in cellular composition caused by hyperoxia and identified the effector cells of the innate immune response, thought to orchestrate pivotal processes resulting in impaired lung development. Next, other systems biology approaches such as spatial transcriptomics will allow the construction of a more dynamic map of events leading to impaired alveolarization. Finally, analyzing the recovery phase after hyperoxia exposure will identify the most severely impaired cell populations unable to recover, which may be responsible for the permanent changes in the lung architecture. In this study, we have described multiple novel aspects of the BPD pathogenesis and identified new pathological pathways as putative drug targets.

## METHODS

### EXPERIMENTAL MODEL AND SUBJECT DETAILS

#### Experimental animals

Pregnant C57BL/6 at embryonic day (E)14 or E17 were purchased from Charles Rivers Laboratories, Saint Constant, QC, Canada. Mice were housed by the Animal Care and Veterinary Service of the University of Ottawa in accordance with institutional guidelines. Newborn mouse pups from dams which delivered on the same day were randomized at day of birth [postnatal day (P) 0] and divided to equal-sized litters of 6 to 8. Following randomization, mice cages were either maintained in room air (normoxia, 21% O_2_) or in normobaric hyperoxia (85% O_2_) from P0 until day of harvest. Hyperoxic environment was maintained in sealed plexiglass chambers with continuous oxygen monitoring (BioSpherix, Redfield, NY). In order to avoid oxygen toxicity and associated confounding factors, nursing dams were rotated between normoxic and hyperoxic group every 48 hours. All mice were maintained in 12/12 hours light/dark cycle and received food *ad libidum*. All developing mice and their nursing dams were euthanized either at P3, P7 or P14 by 10 μl/g intraperitoneal (i.p.) injection of Pentobarbital Sodium (CDMV, Saint-Hyacinthe, QC, Canada). Animals designated for scRNA-seq and FACS analyses received an additional i.p. injection of 10 mU/g heparin sodium (LEO Pharma INc., Thornhill, ON, Canada). All animal procedures were approved by the Animal Care Committee of the University of Ottawa under animal ethics protocol OHRI-1696.

### METHODS DETAILS

#### Sex genotyping of mice

Determination of sex in mouse pups was performed as described previously^66^. Briefly, genomic DNA was isolated from mice tail cuts and regions of interest were amplified by polymerase chain reaction (PCR) in order to determine expression of male-only *Sry* gene, as well as *Il3* gene present in mice of both sexes. Forward and reverse primer sequences are listed in Supplementary table 18. Amplified sequences were visualized by ethidium bromide on 1.5% agarose gel.

#### Mean linear intercept measurement

Pups were euthanized at P3, P7 or P14 by 10 μl/g i.p. injection of pentobarbital sodium (CDMV, Saint-Hyacinthe, QC, Canada). Following euthanasia mice were tracheotomized and lungs were installation-fixed for 5 minutes at 20cm H_2_O hydrostatic pressure with 1.5% (w/v) paraformaldehyde (PFA) (Sigma-Aldrich, Oakville, ON, Canada) and 1.5% (w/v) glutaraldehyde (Sigma-Aldrich, Oakville, ON, Canada) in 150mM HEPES (Sigma-Aldrich, Oakville, ON, Canada) fixation solution with pH 7.4. After isolation, lungs were kept in the fixation solution for 48 hours at 4°C and collected for embedding in paraffin. Paraffin-embedded tissue blocks were sectioned at 4μm and stained with hematoxylin and eosin (H&E) stain. Tissue dehydration, paraffin embedding, sectioning and staining were performed by the University of Ottawa Louis Pelletier Histology Core Facility.

Mean linear intercept (MLI) was estimated in blinded fashion using the Quorum Analysis (Quorum Technologies Inc., Guelph, ON, Canada) software. Briefly, MLI quantification was performed in semi-automated fashion using a 155.34μm line grid moving through sections of interest in defined intervals, where number of intersections between grid line located within alveolar parenchyma and alveolar walls was noted. Average MLI was computed using the formula: 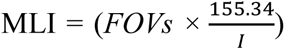, where *FOVs* = fields of view within which intersections were counted, *I* = number of intersections and 155.34 = the length of the grid line. A total of at least 200 *FOVs* were assessed in each lung, corresponding to 10 sections analysed from lungs at P3, 6 sections from lungs at P7 and 4-5 sections from lungs at P14.

#### Stereological estimation of number of alveolar type II cells

The number of AT2 cells was determined in the lungs by stereological principles as described previously^67^. Developing pups were euthanized at P14 by 10 μl/g i.p. injection of pentobarbital sodium (CDMV, Saint-Hyacinthe, QC, Canada), tracheotomized and their lungs were installation-fixed for 5 minutes at 20cm H_2_O hydrostatic pressure with 4% (w/v) paraformaldehyde (PFA) (Sigma-Aldrich, Oakville, ON, Canada), pH 7.4. Following fixation, lungs were stored in PFA solution for 48h at 4°C, embedded *in toto* in 2% agar (Diamed, Mississauga, ON, Canada) and sliced into 2 mm slices. Lung volume was than estimated by the Cavalieri principle exactly as described before^66^. Sliced pieces of each lung were collected for embedding in paraffin. Each paraffin block, containing one pair of lungs from one mouse, was sectioned in agreement with the rules of serial uniform random sampling, where pairs of consecutive, 3μm thick sections, were collected every 200 sections throughout the block. All sections were stained for the AT2 marker Prosurfactant protein C (ProSPC) and quantified using Stereo Investigator® software (MBF Bioscience, Williston, VT, USA). Essentially, AT2 cells were counted in all consecutive sections at 40x magnification by dissector counting. For every pair of lungs, 0.5% of total surface area was analysed. The number of cells was calculated using the following formula: 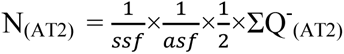, where *ssf* = slide sampling fraction 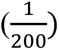, *asf* = area sampling fraction 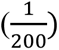 and *ΣQ^-^_(AT2)_* = number of AT2 as counted in both direction in the dissector setting. The number of AT2 cells was normalized to total surface area of the lung. Surface area was calculated as: Sv × N_par_ × V_lung_, where *Sv* = surface density in mm^-1^, *N_par_* = parenchymal fraction and *V_lung_* = lung volume in mm^3^ as estimated by Cavalieri principle. Parenchymal fraction was assessed as described previously. Surface density was assessed stereologically as described before and calculated using following formula: 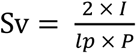, where *Sv* = surface density in mm^-1^, *I* = number of intersections of probe with alveolar surface, *lp* = length of probe/point and *P* = number of points of the probe falling within parenchymal region of the lung^66^. Tissue dehydration, paraffin embedding, sectioning and immunohistochemistry were performed by the University of Ottawa Louis Pelletier Histology Core Facility.

#### Immunohistochemistry

Briefly, 3μm thick paraffin sections were deparaffinized in xylene and rehydrated in decreasing ethanol series. Retrieval was accomplished using an ethylenediaminetetraacetic acid (EDTA) buffer (Bond epitope retrieval solution 2, Leica, Concord, ON, Canada), pH9 for 20 minutes. Sections were stained for proSP-C using an anti-proSP-C antibody (Millipore/Sigma, Entobicoke, ON, Canada) at 1:1500 dilution for 30 minutes followed by detection with horseradish peroxidase (HRP)-conjugated polymer system (Bond Polymer Refine Detection Kit, Leica, Concord, ON, Canada). All sections were then stained using 3,3’-Diaminobenzidine (DAB) as chromogen, counterstained with Hematoxylin and mounted.

#### Fluorescent RNA in situ hybridization

RNA *in situ* hybridization was performed on fresh 4% PFA-fixed paraffin embedded 3μm tissue sections using RNAscope Multiplex Fluorescent Reagent Kit Version 2 (Advanced Cell Diagnostics, Newark, CA, USA) for target detection according to the manual. Firstly, tissue sections were baked for 1 h at 60°C, then deparaffinized and treated with hydrogen peroxide for 10 min at room temperature (RT). Target retrieval was performed for 15 min at 98°C, followed by protease plus treatment for 15 min at 40°C. All probes were hybridized for 2 h at 40°C followed by signal amplification and developing of HRP channels was done according to manual. The following probes were used in the study: 3-Plex negative control probe dapB (#320871), 3-Plex positive control probe_Mm (#320881), Mm-Col1a1-C2 (#319371-C2), Mm-Inhba-C2 (#455871-C2), Mm-Inmt (#486371), Mm-Marco (#510631), Mm-Mrc1-C3 (#437511-C3), Mm-Pecam1 (#316721), Mm-Saa3 (#446841). TSA Plus fluorophores fluorescein Cyanine 3 (1:1500 dilution) and Cyanine 5 (1:3000 dilution) (Perkin Elmer, Waltham, MA, USA) were used for signal detection. Sections were counterstained with DAPI and mounted with ProLong Gold Antifade Mountant (Invitrogen, Carlsbad, CA, USA). Tissue sections were scanned using 3DHISTECH Pannoramic 250 FLASH II digital slide scanner at Genome Biology Unit (Research Programs Unit, Faculty of Medicine, University of Helsinki, Biocenter Finland) using 1 x 40 magnification with extended focus and 7 focus levels. Images were generated using 3DHISTECH Pannoramic 250 FLASH II digital slide scanner at Genome Biology Unit supported by HiLIFE and the Faculty of Medicine, University of Helsinki, and Biocenter Finland.

#### Lung isolation and tissue dissociation

Developing pups designated for single-cell RNA sequencing (scRNA-seq) and Fluorescence activated cell sorting (FACS) analyses were euthanized at P3, P7 or P14 by 10 μl/g i.p. injection of pentobarbital sodium (CDMV, Saint-Hyacinthe, QC, Canada) and received an additional i.p. injection of 10 mU/g (in 10 μl/g volume) heparin sodium (LEO Pharma INc., Thornhill, ON, Canada). Following euthanasia, the chest was opened and the abdominal aorta and vena cava were cut above the liver. The left atrium was perforated and lungs were perfused through the right ventricle with 5 ml of 25 U/ml Heparin Sodium until completely white. Lungs were removed, dissected into individual lobes, and shortly rinsed with Dulbecco’s PBS (DPBS, Lonza, Basel, Switzerland). Dissected lungs were than digested in 5 ml of enzyme mixture at 37°C by gentleMACS™ Octo Dissociator (Miltenyi Biotech, Bergisch Gladbach, Germany). Following costumed dissociation program was used for the digestion: loop 6× (spin 300rpm, 10’’; spin - 300rpm, 10’’); loop 2× (spin 150rpm, 5’’; spin −150rpm, 5’’); loop 2× (spin 20rpm, 5’ 0’’; spin −20rpm, 5’ 0’’); loop 6× (ramp 360rpm, 15’’; ramp −360rpm, 15’’).

The following customized enzymatic mixture was used for lung digestion: i) 2500U Collagenase I (Wothington Biochem., Lakewood, NJ, USA), 30U Neutral Protease (Wothington Biochem., Lakewood, NJ, USA), 500U Deoxyribonuclease (DNAse) I (Sigma-Aldrich, Oakville, ON, Canada); ii) Elastase (Wothington Biochem., Lakewood, NJ, USA), 500U DNAse I; iii) 2500U Collagenase I, 500U DNase I; iv) Collagenase/Dispase (Sigma-Aldrich, Oakville, ON, Canada), 500U DNAse I. All enzyme mixtures were diluted in 5 ml of DPBS supplemented with Mg^2+^/Ca^2+^ (Thermofischer Scientific, Burlington, ON, Canada).

The resulting suspension was filtered through a 100 μm nylon mesh (Termofisher Scientific, Burlington, ON, Canada) and the enzymatic reaction was terminated by 0.9 mM EDTA. The cell suspension was than centrifuged and the resulting pellet was resuspended in 5 ml DPBS (Lonza, Basel, Switzerland), thoroughly filtered through 40 μm filter (Corning Life Sciences, Tewksbury, MA, USA) and centrifuged again. According to its size, the resulting pellet was resuspended in 500 - 1000 μl of cold RBC lysis buffer (Thermofischer Scientific, Burlington, ON, Canada) for 3-5 minutes until white appearance of the suspension was achieved. The cell suspension was than diluted by DPBS (Lonza, Basel, Switzerland) to a total volume of 5ml, centrifuged and washed twice. Cells were counted using both, the Scepter™ automated cell counter (Millipore-Sigma, Burlington, MA, USA) and a manual hematocyter (Bright-Line™ Hematocyter; Sigma-Aldrich, Oakville, ON, Canada).

#### Fluorescent activated cell sorting

The number of cells in single-cell suspension was estimated using a Scepter™ automated cell counter (Millipore-Sigma, Burlington, MA, USA) and total of 0.5×10^6^ cells/sample were resuspended in 200 μl of PBS in 96-well plate. Cells were incubated in the dark with 2 μl/1×10^6^ cells of CD16/32 antibody (Fc block; BD Biosciences, Mississauga, ON, Canada) for 15 minutes at RT. Following blocking, cells were centrifuged and resulting pellets were resuspended in 1:100 mixture of panel of antibodies: FITC-conjugated CD31 (BD Biosciences, Mississauga, ON, Canada), AF647-conjugated CD45 (Southern Biotech, Birmingham, AL, USA), Pe/Cy7-conjugated CD326 (EpCAM; Thermofischer Scientific, Burlington, ON, Canada), Pe-conjugated CD144 (VE-Cadherine; BD Biosciences, Mississauga, ON, Canada). Cells were incubated with antibodies at RT for 30 minutes in dark. Following staining, cells were pelleted by centrifugation and washed 3x with FACS buffer (5% (v/v) FBS and 1mM EDTA in 1×DPBS). All samples were fixed by 4% (w/v) PFA prior to analysis. Flow cytometry was performed using a BD LSR Fortessa (Beckton Dickinson Biosciences, Franklin Lakes, NJ, USA) at the Ottawa Hospital Research Institute (OHRI) core facility. Sample compensation was performed using BD FACSDIVA software and data analysis were performed with FlowJo v10 software (FlowJo LLC, Ashland, OR, USA).

#### Multiplexing individual samples for scRNA-seq

Multiplexing was performed according to the MULTI-seq protocol^6^. Following the preparation of single-cell suspension, cells were counted and a total of 0.5×10^6^ cells/sample were resuspended and pelleted in a 96-well plate at 400 × g for 5 minutes. The resulting pellet was resuspended in 150 μl of 200 nM anchor/200 nM barcode solution (kindly provided by Prof. Zev Gartner from University of California, San Francisco). The lipid-modified DNA oligonucleotide (LMO) anchor and a unique “sample barcode” oligonucleotides were added to each sample in order to be multiplexed, with each sample receiving a different sample barcode (Supplementary Figure 1a). Samples were then incubated for 10 minutes at room temperature (RT) (Supplementary Figure 1a). After 10 minutes, samples were supplemented with 200 nM common lipid-modified co-anchor to stabilize the membrane residence of barcodes. Samples were incubated on ice for additional 5 minutes and pelleted at 400 × g for 5 minutes. Barcode-containing media was then removed, and the resulting cell pellet was washed twice with 1% FBS (Sigma-Aldrich, Oakville, ON, Canada) in 1× DPBS (Lonza, Basel, Switzerland). After the final wash, cells were resuspended in 1× DPBS + 1% FBS, counted and samples were pooled together at 1:1 ratio while maintaining the final concentration of 500-1000 cells/μl. Viability and cell counts were assessed using a manual hematocyter (Bright-Line™ Hematocyter; Sigma-Aldrich, Oakville, ON, Canada), and only samples with viability ≥ 80% were further processed by 10× Chromium.

#### ScRNA-seq library preparation and sequencing

Single-cell suspensions were processed using the 10x Genomics Single Cell 3’ v3 RNA-seq kit. Gene expression libraries were prepared according to the manufacturer’s protocol. MULTI-seq barcode libraries were retrieved from the samples and libraries were prepared independently, as described previously^6^. Final libraries were sequenced using NextSeq500 (Illumina). Expression libraries were sequenced in order for time course libraries to reach an approximate depth of 20,000-25,000 reads/ cell.

### DATA ANALYSES, QUANTIFICATION AND STATISTICAL ANALYSES

#### Processing of raw sequencing reads

Raw sequencing reads from the gene expression libraries were processed using CellRanger v3.0.2, aligning reads to the mm10 build of the mouse genome. Except for explicitly setting --expect-cells=25000, default parameters were used for all samples. MULTI-seq barcode libraries were simply trimmed to 28bp using Trimmomatic^18^ (v0.36) prior to demultiplexing.

#### Demultiplexing expression data with MULTI-seq barcode libraries

Demultiplexing was performed using the deMULTIplex R package (v1.0.2) (https://github.com/chris-mcginnis-ucsf/MULTI-seq). The key concepts for demultiplexing are described in McGinnis *et al*^6^. Briefly, the tool considers the barcode sequencing reads and counts the frequency with which each of the sample barcodes appears in each cell. Then, for each barcode, the distribution of counts in cells is assessed and an optimal quantile threshold to deem a cell positive for a given barcode is determined. Cells positive for more than one barcode are classified as doublets and removed. Only cells positive for a single barcode are retained for further analysis (Supplementary Figure 1b). As each barcode corresponds to a specific sample in the experiment, the sample annotations can then be added to all cells in the data set.

#### Data quality control and processing

Quality control was first performed independently on each 10x Genomic library, and all main processing steps were performed with Seurat v3.0.2^68^. Expression matrices for each sample were loaded into R as Seurat objects, retaining only cells, in which more than 200 genes were detected. Cells with a high percentage of mitochondrial gene expression were removed. Expression values were normalized with standard library size scaling and log-transformation. The top 2000 variable genes were detected in Seurat using the “vst” selection method. Expression values were scaled and the following technical factors were regressed out: percentage of mitochondrial reads, number of RNA molecules detected, and cell cycle scores. PCA was performed on the highly variable genes and UMAP embeddings were calculated from the first 30 principal components. Differential expression between normal and impaired populations in the scRNA-seq data was performed using the Wilcoxon Rank Sum test implemented by the FindMarkers function in Seurat. RNA velocity analysis was performed by first generating spliced and unspliced transcript count matrices using Velocyto^69^. Velocity estimates were calculated using the scVelo^70^ python package with default parameters.

#### Statistical analysis

Data are presented as means ± SD. All statistical analyses were performed with GraphPad Prism 8.0. The presence of potential statistical outliers was determined by Grubbs’ test. Differences in case of two-member groups were evaluated either by unpaired Student’s *t*-test, or multiple Student’s *t*-test with correction for multiple comparisons using Holm-Sidak method. *P* values < 0.05 were considered as significant and depicted as following: *P* values < 0.05: *; *P* values < 0.01: **; *P* values < 0.001: ***; *P* values < 0.0001: ****.

## AUTHORS CONTRIBUTIONS

KM-H and M-I conceived and directed the study, performed *in vivo* experiements, cell isolations and scRNA-seq analyses and wrote the manuscript. C-D performed scRNA-seq analyses and edited the manuscript. CD-C performed FACS and stereological analyses. L-F performed stereological analyses. A-N performed fluorescent RNA *in situ* hybridization. H-E analyzed scRNA-seq data. R-L analyzed data from *in vivo* studies. J-RP, H-M, V-BC and T-B supervised experiments and provided essential equipment and infrastructure.

## ACKNOWLEDGEMENTS

The authors acknowledge the assistance of Martin Kang, PhD. (Ottawa Hospital Research Institute) with interpretation of scRNA-seq analyses of epithelial cells, Shumei Zhong (Ottawa Hospital Research Institute) and Sharlene Faulkes (University of Ottawa Louis Pelletier Histology Core Facility) for expert technical assistance and Prof. Zev Gartner (University of California, San Francisco) for kindly providing barcodes for multiplex labelling. Authors also acknowledge Michaela Kmaková for assistance with graphical design. This study was supported by the Canadian Institutes of Health Research (CIHR), the Finnish Sigrid Juselius Foundation, the Finnish Foundation for Pediatric Research, the German Research Foundation (Deutsche Forschungsgemeinschaft), the Ontario Institute for Regenerative Medicine (OIRM), the Stem Cell Network Heart and Stroke Foundation Canada, the Ontario Graduate Scholarship, the Lung Association Breathing as One, the Molly Towel Perinatal Research Foundation, the European Respiratory Society, Société Française de Neonatologie and the Canada Foundation for Innovation John R. Evans Leaders Fund.

---

**Supplementary figure 1.**
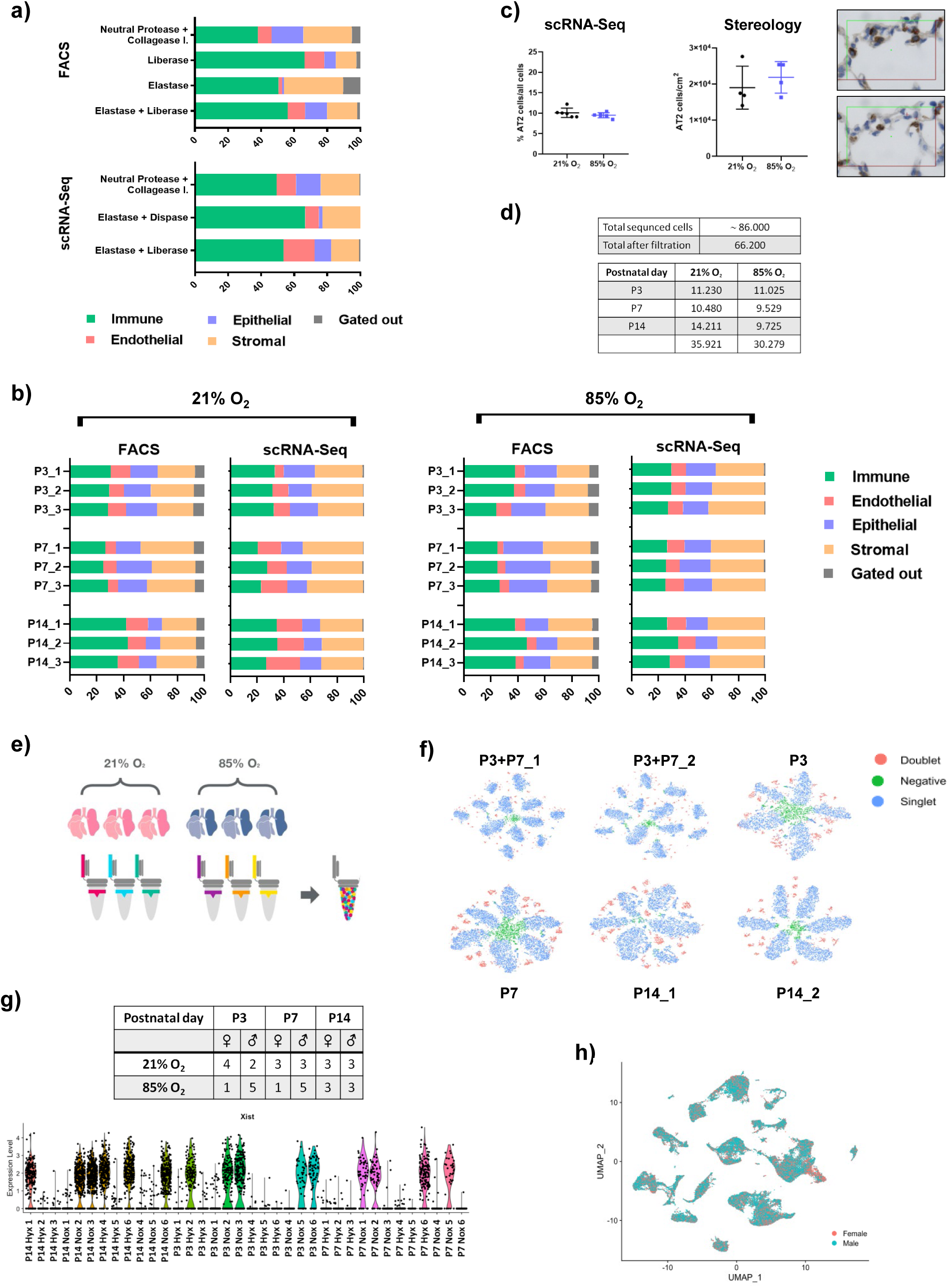
Optimization of single-cell isolation protocol. a) Relative contribution of immune, epithelial, endothelial and stromal cells after digestion with various enzyme buffers as assessed by FACS and scRNA-seq analysis. n = 3/group. b) Comparison of the relative contribution of immune, epithelial, endothelial and stromal cells after single-cell isolation between individual lungs isolated at P3, P7 and P14 as assessed by FACS and scRNA-seq analysis. c) Relative proportion of alveolar epithelial type 2 (AT2) cells in normally and aberrantly developing lungs at P14 assessed by scRNA-seq and by stereology. Representative lung sections illustrate AT2-specific Pro-SPC staining and counting frame used for stereological assessment. Data are presented as means ± SD. Statistical analyses were performed with GraphPad Prism 8.0. The presence of potential statistical outliers was determined by Grubbs’ test. Significance was evaluated by unpaired Student’s *t-* test. n =4-6/group. d) Distribution of analyzed cells across individual experimental groups. e) Single cell suspensions from individual lungs were multiplex-labeled and pooled together prior to scRNA-seq in order to eliminate batch variability. f) tSNE plots from independent sets of scRNA-seq demonstrating the multiplex bar-labeling of individual samples within each batch. n = 6 or 12/experiment. g) Sex distribution as determined by PCR and expression of *Xist* gene. h) uMap representing the sex distribution in all 22 main identified clusters.

**Supplementary figure 2.**
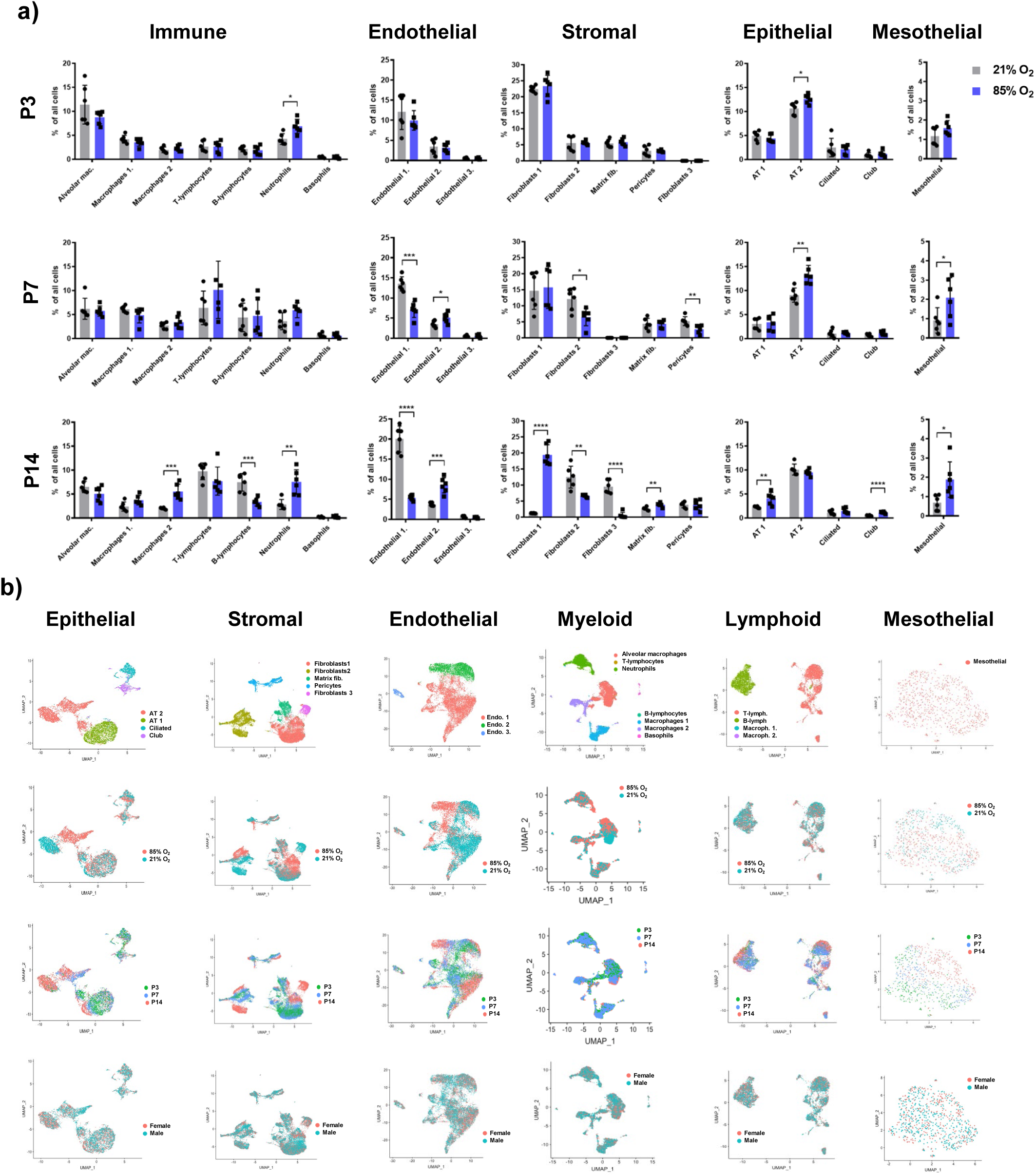
Single-cell isolation and multiplex labeling prior to scRNA-seq. a) The relative contribution of individual clusters within immune, endothelial, mesenchymal, epithelial and mesothelial cells in normal and aberrantly developing lungs at P3, P7 and P14. Data are presented as means ± SD. Statistical analyses were performed with GraphPad Prism 8.0. The presence of potential statistical outliers was determined by Grubbs’ test. Significance was evaluated by unpaired Student’s *t-* test. P values < 0.05: *; P values < 0.01: **; P values < 0.001: ***; P values < 0.0001: ****; n =6/group. b) uMaps representing distribution of epithelial, mesenchymal, endothelial, myeloid, lymphoid and mesothelial cells based on cluster identity from main 22 clusters, oxygen experimental conditions, timepoint and sex.

**Supplementary figure 3.**
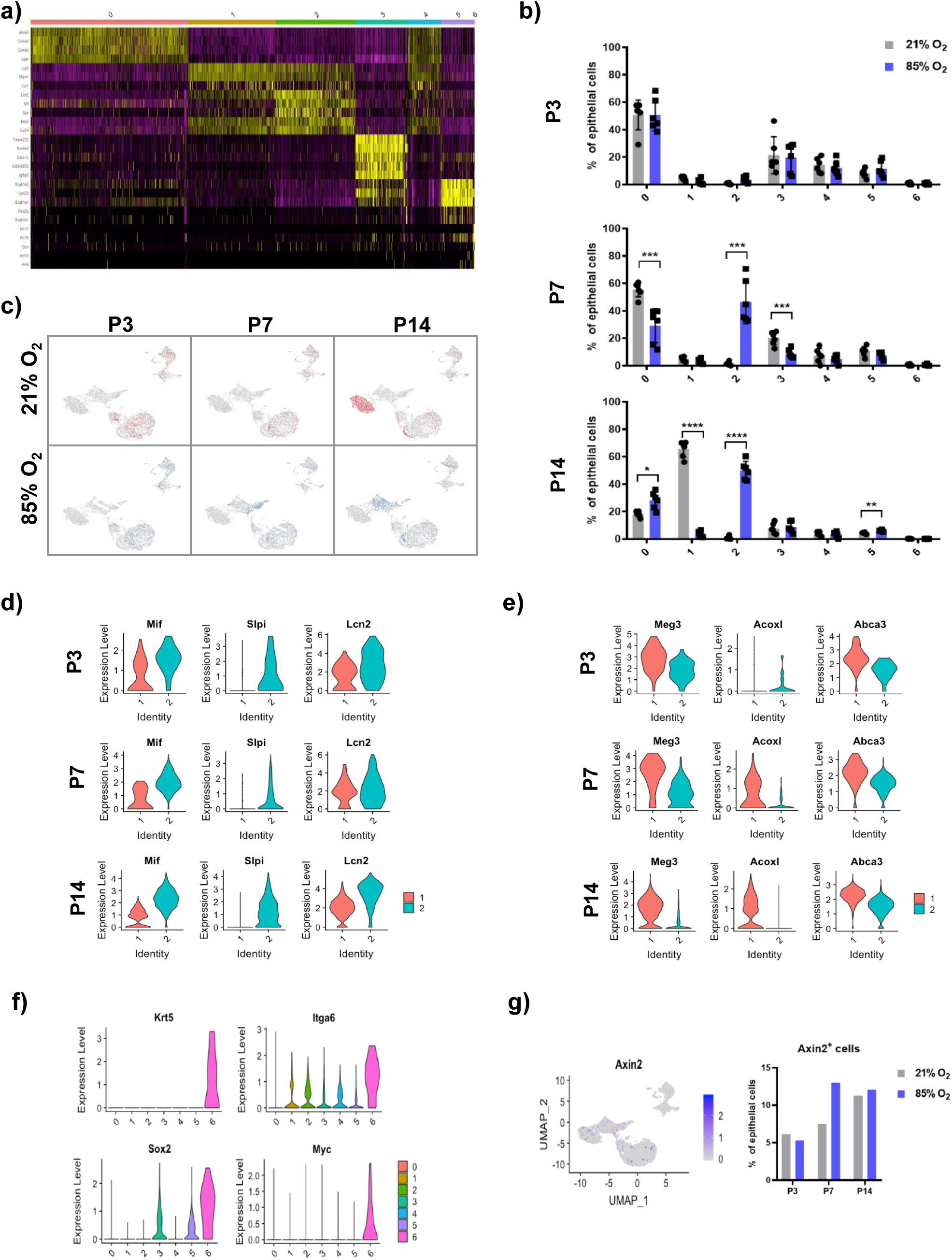
Cellular composition of normal and aberrantly developing lung epithelium. a) Heatmap of the most differentially expressed genes across the 7 epithelial clusters. b) Hyperoxia exposure altered gene expression in developing lung epithelium at all investigated timepoints. Data are presented as means ± SD. Statistical analyses were performed with GraphPad Prism 8.0. The presence of potential statistical outliers was determined by Grubbs’ test. Significance was evaluated by unpaired Student’s *t-* test. P values < 0.05: *; P values < 0.01: **; P values < 0.001: ***; P values < 0.0001: ****. n =6/group. c) uMaps representing the temporal changes in gene expression in alveolar epithelial (AT) cells during normal and aberrant late lung development. d) Violin plots depicting AT2_2 cluster (cluster 2) specific genes. e) Violin plots depicting AT2_1 cluster (cluster 1) specific genes. f) Violin plots depicting main identifiers of distal alveolar stem cells population in developing lung epithelium. g) Relative contribution of *Axin2*^+^ cells in developing lung epithelium at P3, P7 and P14.

**Supplementary figure 4.**
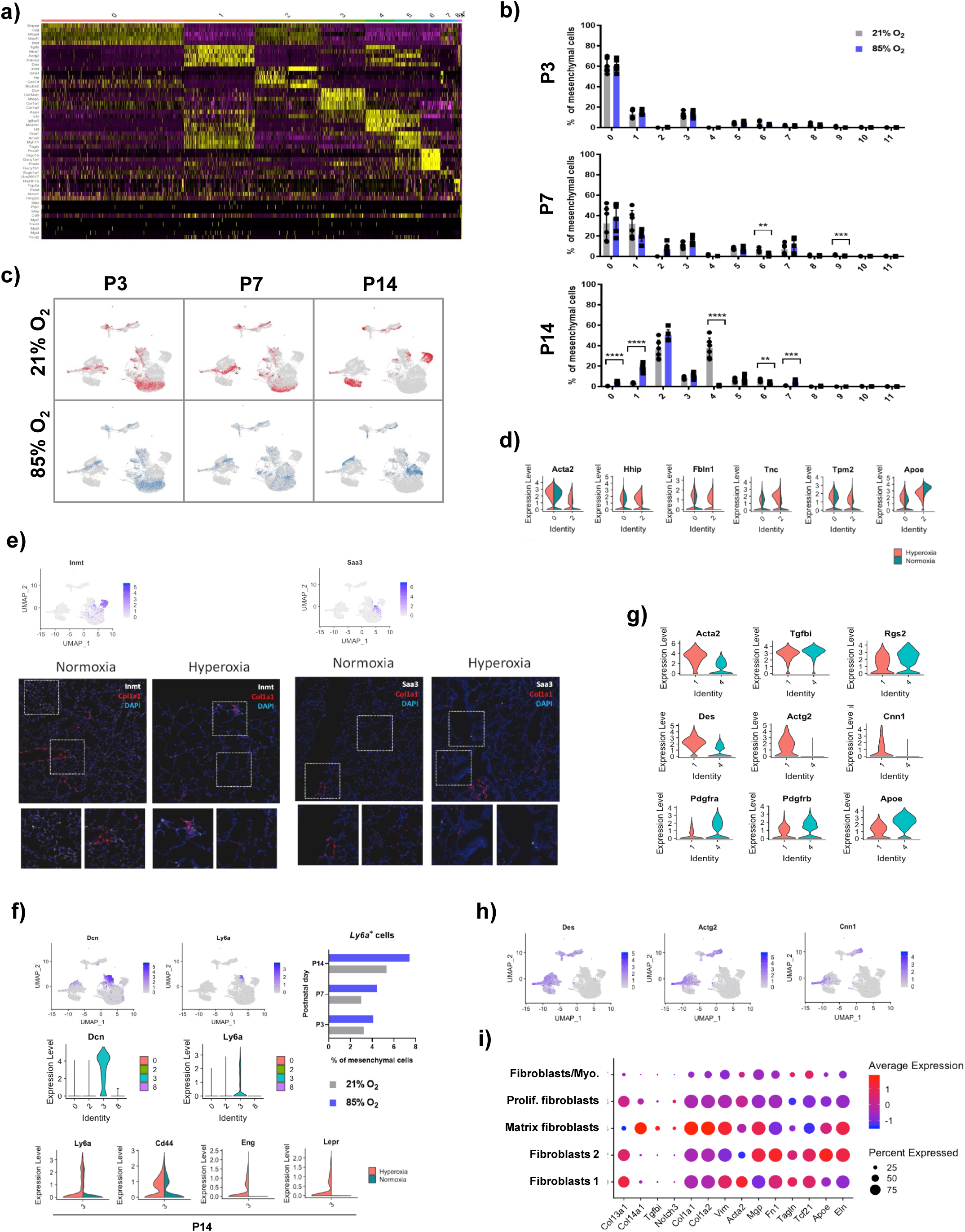
Cellular composition of normal and aberrantly developing lung mesenchyme. a) Heatmap of the most differentially expressed genes across the 12 mesenchymal clusters. b) Hyperoxia exposure altered gene expression in developing lung mesenchyme at all investigated timepoints. Data are presented as means ± SD. Statistical analyses were performed with GraphPad Prism 8.0. The presence of potential statistical outliers was determined by Grubbs’ test. Significance was evaluated by unpaired Student’s *t-* test. P values < 0.01: **; P values < 0.001: ***; P values < 0.0001: ****. n =6/group. c) uMaps representing the temporal changes in gene expression in lung mesenchyme during normal and aberrant late lung development. d) Violin plots depicting Fibroblasts 1 cluster (cluster 0) and Fibroblasts 2 cluster (cluster 2)-specific gene expression patterns. e) uMaps and representative images from fluorescent RNA *in situ* hybridization, for top identifiers of normoxic and hyperoxic fractions of Fibroblast 2 cluster. Normoxia-specific marker Inmt and hyperoxia-specific marker *Saa3* (white) and *Col1a1* (red). Magnification: 40x. f) uMaps and violin plots depicting specific localization of *Ly6a*^+^ cells within Matrix fibroblasts cluster (cluster 3). Number of *Ly6a*^+^ cells and expression of stem cell-markers were specifically increased in developing lungs by exposure to hyperoxia. n =6/group. g) Violin plots depicting SMCs (cluster 1) and SMC/Myo (cluster 4)-specific gene expression patterns. h) uMaps depicting the expression of markers of mature SMCs: *Des*, *Actg2* and *Cnn1*. i) Dotplot depicting expression pattern of fibroblast-associated markers across different fibroblast populations in developing lung.

**Supplementary figure 5.**
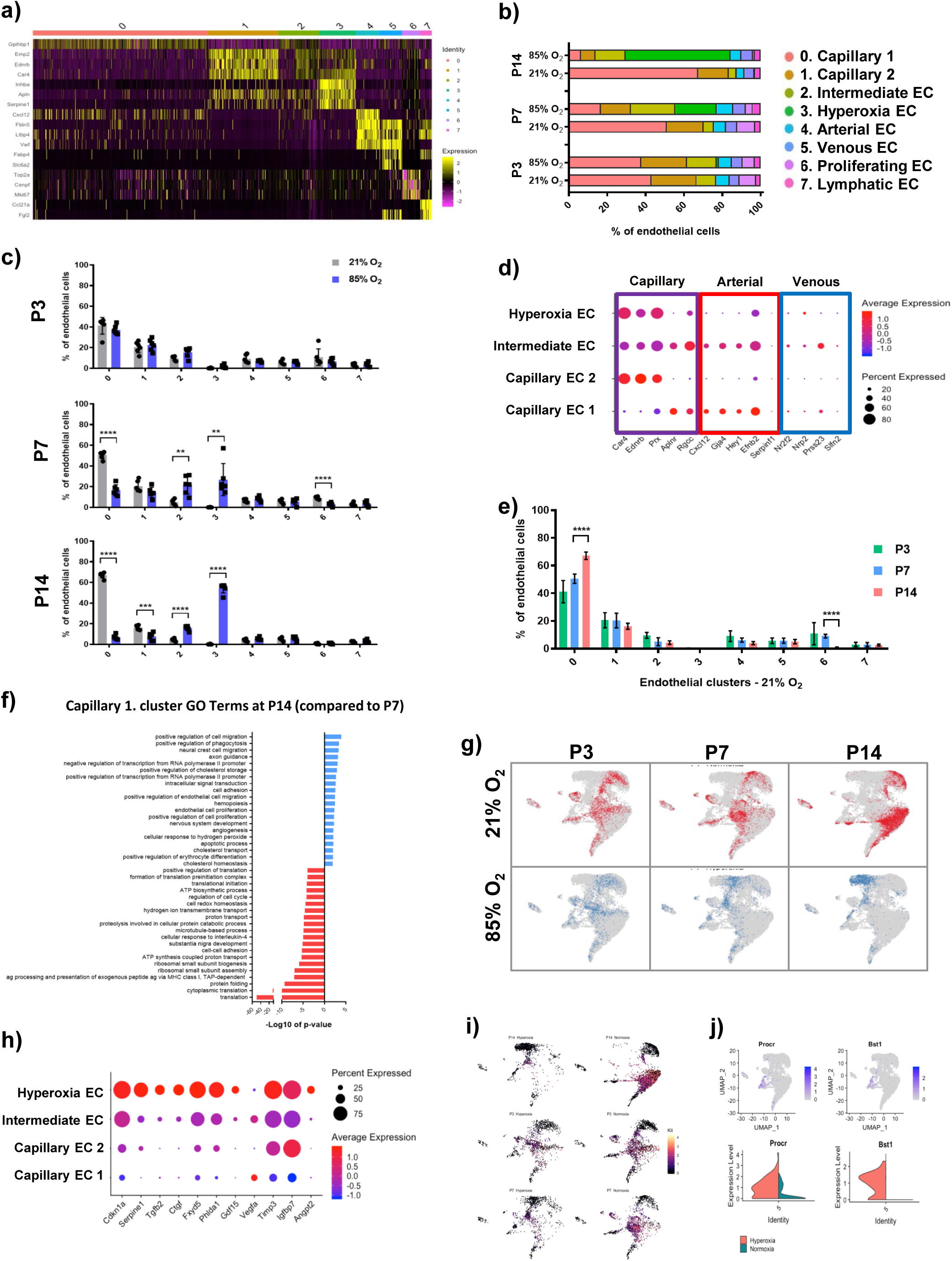
Cellular composition of normally and aberrantly developing lung endothelium. a) Heatmap of the most differentially expressed genes across the 8 endothelial clusters. b) Relative contribution of individual endothelial clusters changed significantly during the development and after exposure to hyperoxia. n = 6/group. c) Hyperoxia exposure altered gene expression in developing lung endothelium at all investigated timepoints. Data are presented as means ± SD. Statistical analyses were performed with GraphPad Prism 8.0. The presence of potential statistical outliers was determined by Grubbs’ test. Significance was evaluated by unpaired Student’s *t-* test. P values < 0.01: **; P values < 0.001: ***; P values < 0.0001: ****. n =6/group. d) Dotplot displaying expression patterns specific to diverse populations within capillary endothelium. e) Temporal changes in relative contribution of individual endothelial populations in normally developing lung. Data are presented as means ± SD. Statistical analyses were performed with GraphPad Prism 8.0. The presence of potential statistical outliers was determined by Grubbs’ test. Significance was evaluated by unpaired Student’s *t-* test. P values < 0.0001: ****. n =6/group. f) Top 20 GoTerm pathways differentially regulated in Cluster 1 at P14 compared to P7. g) uMaps representing the temporal changes in gene expression in lung endothelium during normal and aberrant late lung development. h) Dotplot displaying expression patterns across capillary endothelium populations. i) uMaps depicting expression of *Kit* in normally and aberrantly developing lungs. j) uMaps and violin plots depicting cluster 5-specific expression of *Procr* and *Bst1* in normally and aberrantly developing lungs.

**Supplementary figure 6.**
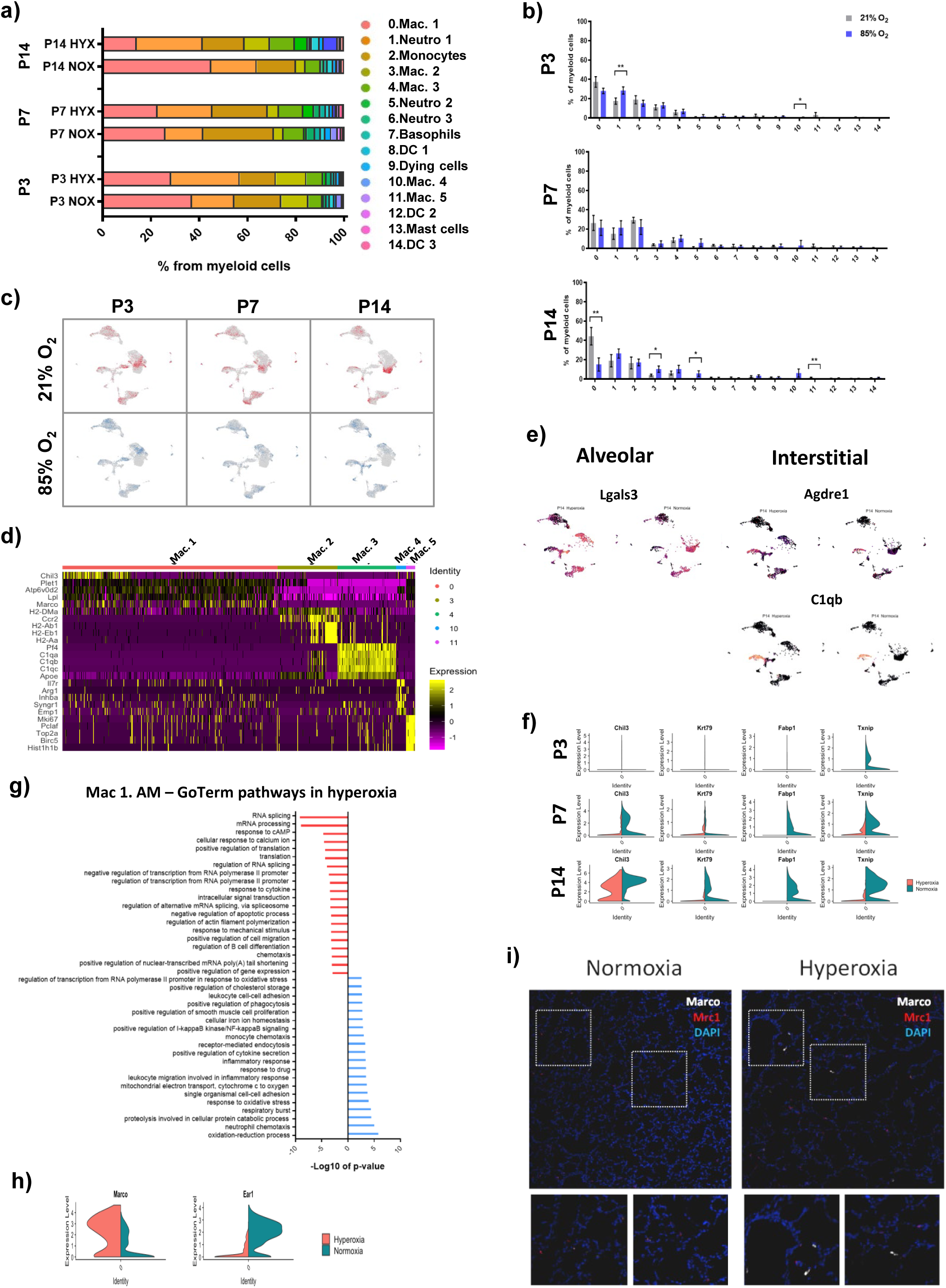
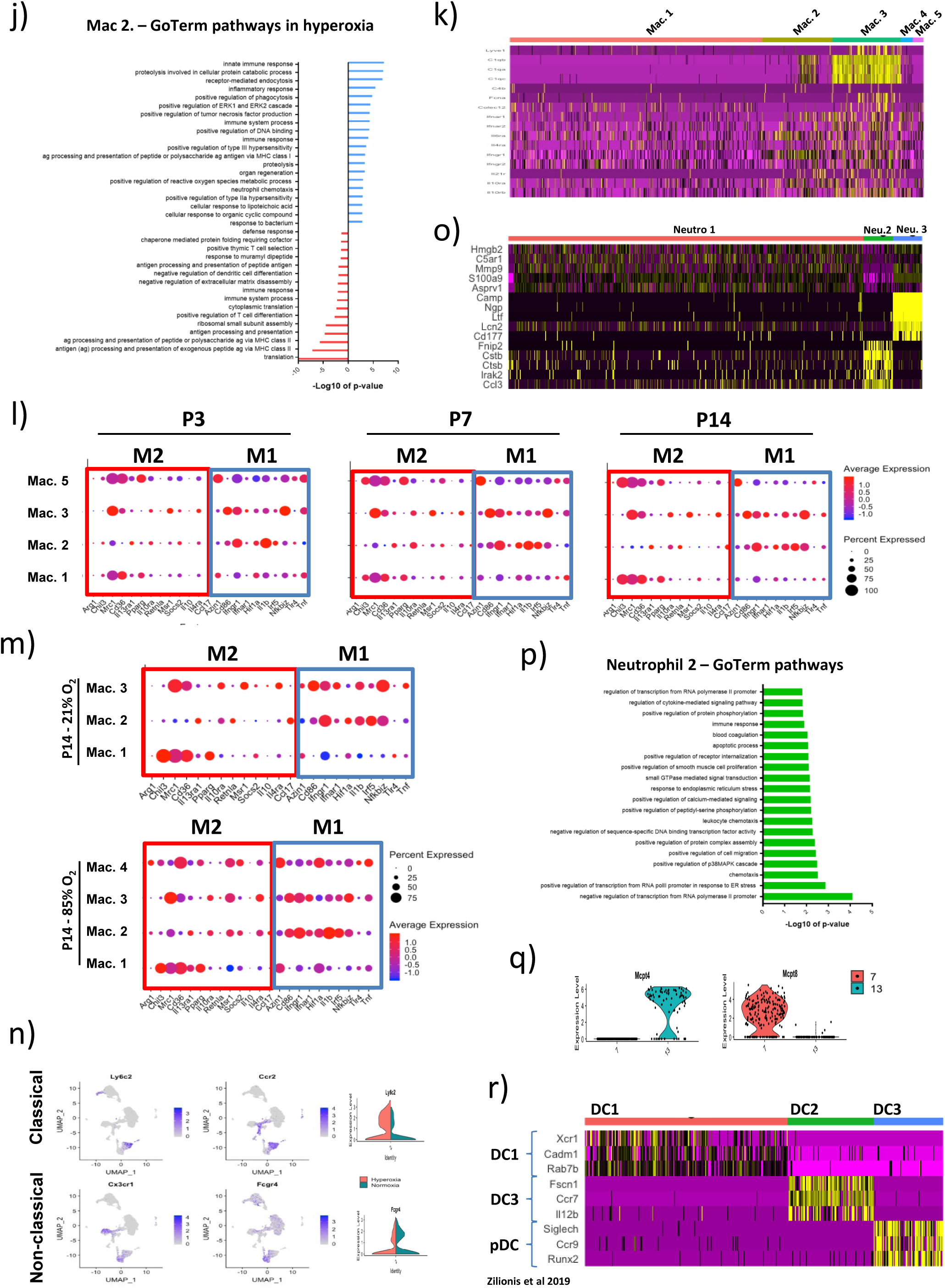
Cellular composition of normal and aberrantly developing lung myeloid populations. a) Relative contribution of individual myeloid clusters changed significantly during the development and after hyperoxia exposure. n = 6/group. b) Hyperoxia exposure altered gene expression in developing myeloid populations at all investigated timepoints. Data are presented as means ± SD. Statistical analyses were performed with GraphPad Prism 8.0. The presence of potential statistical outliers was determined by Grubbs’ test. Significance was evaluated by unpaired Student’s *t-* test. P values < 0.05: *; P values < 0.01: **. n =6/group. c) uMaps representing the temporal changes in gene expression in myeloid populations during normal and aberrant lung development. d) Heatmap of most differentially expressed genes in five distinct macrophage clusters. e) uMaps depicting expression of markers of alveolar and interstitial macrophage populations in normally and aberrantly developing lungs. f) Violin plots depicting differential gene expression patterns in alveolar macrophages induced in lungs by hyperoxia. g) Top 20 GoTerm pathways impacted by hyperoxia in Macrophage 1 cluster. h) Violin plots depicting the oxygen-specific expression of *Marco* and *Ear1* in alveolar macrophages. i) Representative images from fluorescent RNA *in situ* hybridization for hyperoxia-specific marker *Marco* (white) and macrophage marker *Mrc1* (CD206; red). Magnification: 40x. j) Top 20 GoTerm pathways impacted by hyperoxia in Macrophage 2 cluster. k) Heatmap depicting the Macrophage 3 cluster-specific expression of inflammatory mediators. l) Dotplot displaying expression patterns of M1 and M2 macrophages-specific markers across four macrophage populations identified in our study. m) Dotplot displaying hyperoxia-induced changes in expression patterns of M1 and M2 macrophages-specific markers across macrophage populations identified in our study. n) uMaps and violin plots showing the expression patterns of classical and non-classical monocytes markers. o) Heatmap of most differentially expressed genes in three distinct neutrophil clusters. p) Top 20 GoTerm pathways induced in Neutrophils 2 cluster. q) Violin plots showing the cluster-specific expression of top identifiers of basophils (cluster 7) and mast cells (cluster 13). r) Heatmap of most differentially expressed genes in three cluster of dendritic cells identified in our study and their relation to dendritic clusters described by *Zilinois et al*.^60^.

